# The emergence and molecular evolution of H5N1 influenza viruses in United States dairy cattle

**DOI:** 10.64898/2026.03.30.713641

**Authors:** Jonathan E. Pekar, Karthik Gangavarapu, Alvin Crespo-Bellido, Thomas P. Peacock, Joel O. Wertheim, Gytis Dudas, Jeffrey B. Joy, Meera Chand, Florence Débarre, Praneeth Gangavarapu, Daniel H. Goldhill, Natalie Groves, Xiang Ji, Lorena Malpica Serrano, Louise Moncla, Angela L. Rasmussen, Christopher Ruis, Divya Venkatesh, Moritz U. G. Kraemer, Oliver G. Pybus, Kristian G. Andersen, Marc A. Suchard, Martha I. Nelson, Philippe Lemey, Michael Worobey, Andrew Rambaut

## Abstract

Prior to 2024, highly pathogenic avian influenza H5N1 clade 2.3.4.4b viruses circulated predominantly in wild birds and poultry. In 2024 and 2025, 2.3.4.4b genotypes B3.13 and D1.1 were detected in United States dairy cattle. Using whole-genome and segment-specific phylodynamic inference, we estimate that B3.13 and D1.1 spilled over from wild birds into dairy cattle in late 2023 and late 2024, respectively. Spillover occurred shortly after the formation of the reassortant genotypes and was followed by months of cryptic transmission prior to detection. We found that both B3.13 and D1.1 evolved at higher rates in cattle relative to birds, primarily due to relaxed purifying selection. Site-specific analyses identified genomic sites under positive selection in cattle relative to birds, indicating adaptation and likely contributing to improved viral fitness after spillover. Intensified genomic surveillance in dairy cattle is essential as population immunity introduces additional selection pressures, with ever-changing risk for human emergence.

## Introduction

In March 2024 a series of avian influenza virus (AIV) cases were detected in United States (US) dairy cattle^1–3^, a species not previously recognized as capable of sustaining AIV transmission. These cases resulted from spillover from birds to cattle of an H5N1 clade 2.3.4.4b genotype B3.13 virus. Early transmission was concentrated in Texas, suggesting the state acted as the epicenter of the outbreak, and the first human infection associated with the outbreak was identified in March 2024 in a dairy worker in Texas^4^. By May 2024, most reported cases and sequenced genomes originated from outside of Texas. By the end of 2024, 917 US dairy premises had documented cases, and the virus had diversified into seven major subclades^5^. Transmission continued into 2025, establishing H5N1 transmission in a novel livestock host at unprecedented geographic scale.

In January 2025, a second clade 2.3.4.4b genotype (D1.1) was detected in dairy cattle in Nevada and shortly thereafter in Arizona, through the US Department of Agriculture’s National Milk Testing Strategy highly pathogenic avian influenza (HPAI) surveillance program^6^. By the time of its detection in cattle, D1.1 had already become widespread in North American wild birds and has since been a major source of poultry outbreaks, as well as severe human infections, including a fatal case in Louisiana^7^ and a hospitalization in British Columbia^8^. In February 2025, Nevada reported its first zoonotic case of H5N1 in an individual exposed to infected dairy cattle^9^. The outbreaks in Nevada and Arizona affected at least 11 and 5 premises, respectively. In contrast to B3.13, these two outbreaks appeared genetically distinct, raising the possibility of separate introductions into dairy herds.

Since dairy cattle are a newly recorded host species for H5N1, the evolutionary dynamics of the virus in this host remain poorly understood. Sustained circulation in cattle could reflect adaptation to novel tissues and/or transmission environments. Transmission among cattle is thought to occur primarily through the mammary gland rather than the respiratory tract, although this remains unresolved^10^ and may influence the selective pressures acting on the virus. Here we use phylogenetic approaches to analyze the emergence, evolution, and natural selection of B3.13 and D1.1 viruses in US dairy cattle and closely related avian viruses. We demonstrate that both lineages emerged in dairy cattle shortly after their emergence in wild birds, with evidence for several months of cryptic circulation of each lineage in cattle before their discovery. We find that, despite differences in the extent of circulation in dairy cattle, both the B3.13 and D1.1 lineages experienced a relaxation of selection once circulating in dairy cattle. Finally, by identifying sites under positive selection in avian and bovine hosts, we identify regions of the viral genome that experience altered selective pressures following host shifts and which may represent foci of host-specific adaptation.

## Results

### Timing of the emergence of B3.13 in dairy cattle and wild birds

We first focused on the emergence of the B3.13 lineage responsible for the initial H5N1 outbreak in US dairy cattle (Fig. 1). To confirm that the B3.13 US dairy cattle outbreak resulted from a single introduction from wild birds, we inferred separate maximum likelihood phylogenies for each genome segment. We find that the viruses sampled from dairy cattle form a monophyletic clade in each segment (Fig. S2), consistent with a single introduction of H5N1 into cattle and in line with previous results^11^. As the PA, HA, NA, and MP segments of B3.13 are all derived from the A1 genotype (Fig. S1), we concatenated these four segments to increase phylogenetic resolution (A1 concatenated alignment). The dairy cattle outbreak clade was again monophyletic on the tree inferred from the concatenated alignment, corroborating the single-segment results (Fig. 1a, Fig. S2). The basal B3.13 clade structure is concordant with a tree inferred from concatenated whole genomes of B3.13 viruses, with the earliest human B3.13 sequence (A/Texas/37/2024(H5N1)) forming a sibling lineage to the cattle outbreak clade while remaining more closely related to it than the only other B3.13 genomes—four sequences sampled from wild animals—which appear distinct from this clade (Fig. S3).

**Figure 1.**
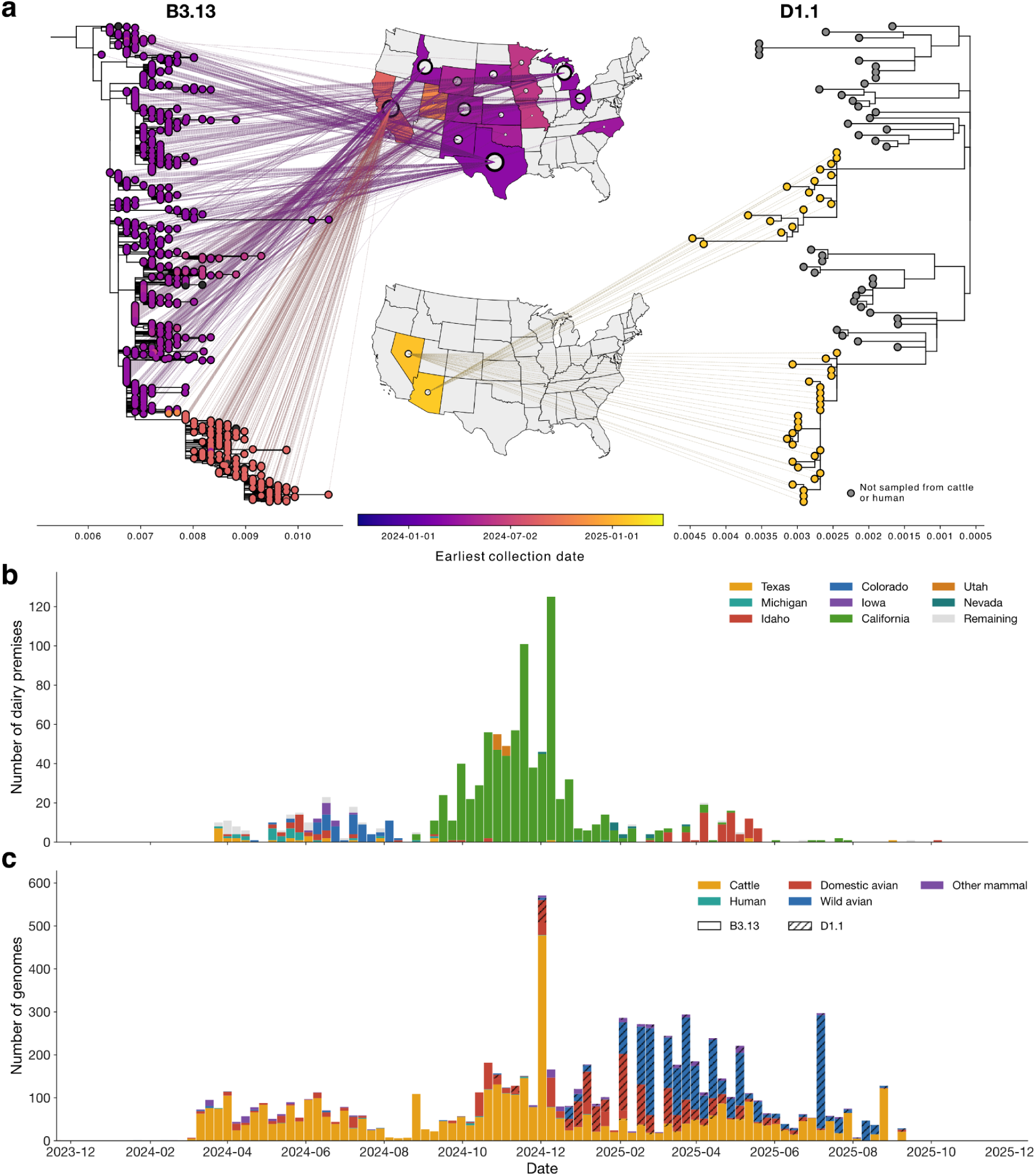
Emergence of B3.13 and D1.1 in US dairy cattle. **a**, Maximum likelihood phylogenies of B3.13 (left) and D1.1 (right). Lines connect dairy cattle and human virus genomes to the US states in which they were sampled. US states are colored by the earliest collection date of genomes from dairy cattle or humans in that state. Only tree tips corresponding to dairy cattle and human viruses are colored by the state of sampling; tips and corresponding states are colored by collection date for genomes with metadata and by publication date for genomes lacking collection dates. (Fully annotated phylogenies of B3.13 and D1.1 are provided in Figures S3i and S7, respectively). **b**, Number of dairy premises with reported H5N1 infections per week in the US. States with <10 affected premises are grouped into a single “Remaining” category. **c**, Number of sequenced B3.13 and D1.1 genomes sampled per week in the US. Dates correspond to collection dates where available (n=4093) and publication dates otherwise (n=3945).

To determine when B3.13 emerged in dairy cattle, we performed a Bayesian phylodynamic analysis of H5N1 viruses sampled from 2021 to 2024 using the A1 concatenated alignment. Because evolutionary rates can vary among host species for the same pathogen, including for AIVs^12^, accommodating such rate differences is needed to accurately estimate phylogenies and divergence times. We therefore allowed the evolutionary rate of the dairy cattle outbreak to differ from that of the background avian viruses. We inferred a time of the most recent common ancestor (tMRCA) of the dairy cattle outbreak clade of mid-December 2023 (95% highest posterior density interval [HPDI]: 9 November 2023–6 January 2024; Fig. 2a–b), suggesting that H5N1 was circulating cryptically for at least 2–3 months prior to detection of the initial infections on dairy farms in Texas. The inferred time of the parent of the MRCA (tPMRCA)—representing the earliest plausible time for the spillover of B3.13 into dairy cattle—was late October 2023 (95% HPDI: 30 September 2023–22 November 2023).

**Figure 2.**
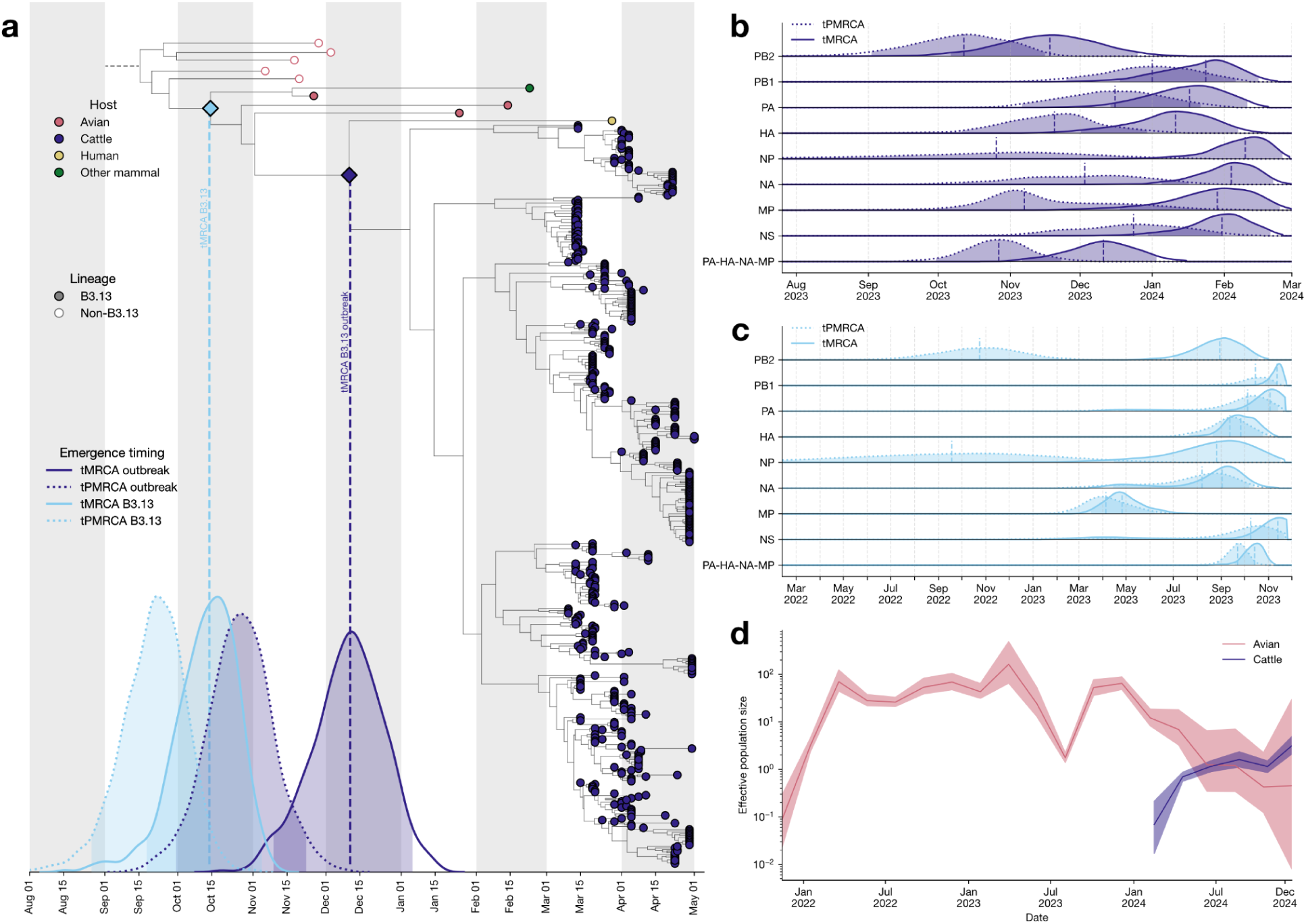
Emergence of B3.13. **a**, Time-calibrated phylogeny inferred from a concatenated alignment of the PA, HA, NA, and MP segments, including B3.13 dairy cattle and human viruses and background avian influenza viruses from the A1 genotype (see Methods). Although B3.13 is defined at the whole-genome level, the four concatenated segments fall within the A1 genotype. Tips are colored by host species, and lineage membership is indicated by filled versus open points. The time to the most recent common ancestor (tMRCA) of the dairy cattle outbreak is shown by the purple density distribution, with a solid line; the time to the parent MRCA (tPMRCA) is shown by the purple density with a dotted line; the vertical dashed line indicates the median tMRCA, and the purple diamond marks the corresponding node in the phylogeny. The overall tMRCA of B3.13 is shown by the light blue density with a solid line; the tPMRCA is shown by the light blue density with a dotted line; the vertical light blue dashed line indicates the posterior median tMRCA, and the light blue diamond marks the corresponding node. See Figure S5 for the complete phylogeny. **b**, Segment-specific and concatenated posterior density estimates of the tMRCA and tPMRCA of B3.13. **c**, Segment-specific and concatenated posterior density estimates of the tMRCA and tPMRCA of the B3.13 dairy cattle outbreak. **d**, Effective population size trajectories for the A1 genotype in avian and cattle hosts. Lines indicate posterior median effective population size; shaded regions indicate the 95% highest posterior density interval.

**Figure 3.**
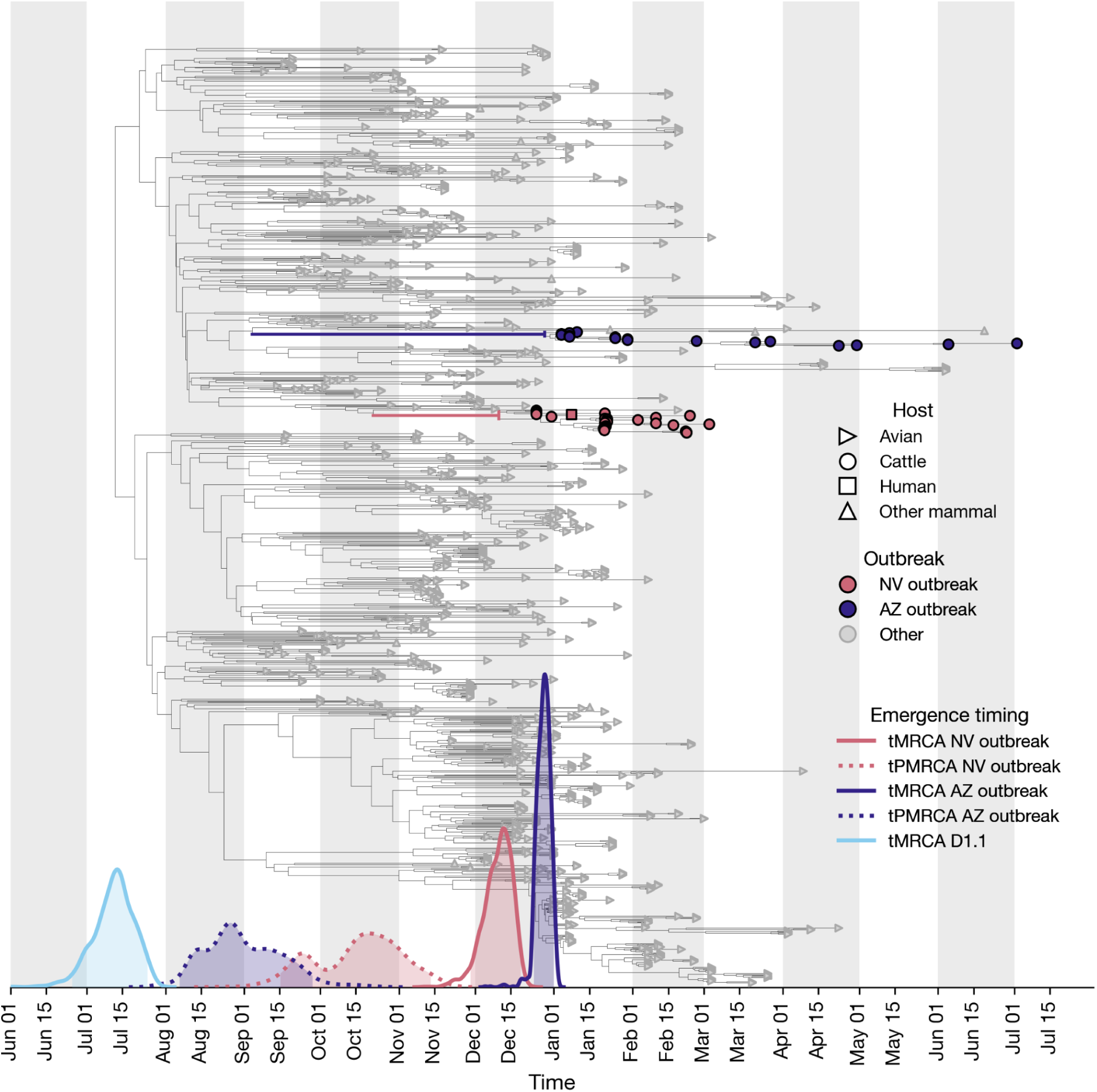
Emergence of D1.1. Time-calibrated phylogeny of D1.1, with node shape indicating host and node color indicating which dairy cattle outbreak the tree tips correspond to. Posterior density plots show the tMRCA (purple) and tPMRCA (light red) of the two dairy cattle outbreaks, as well as the overall tMRCA of D1.1 (light blue). Branches connecting the PMRCA and MRCA of each dairy cattle outbreak are highlighted and colored consistently with the corresponding tips and density plots.

We also performed equivalent Bayesian phylodynamic analyses for each individual influenza gene segment (Fig. S1; see Methods). Estimates were broadly consistent with those from the concatenated alignment (Fig. 2b), albeit with greater uncertainty, as expected for a subgenomic analysis (Fig. S4).

The inferred tMRCA of all B3.13 viruses was mid-October (95% HPDI: 18 September 2023–5 November 2023), only a few weeks earlier than when B3.13 was likely introduced into dairy cattle. The tPMRCA of the B3.13 lineage as a whole was mid-September (95% HPDI: 26 August 2023–14 October 2023), suggesting that B3.13 spilled over into dairy cattle shortly after its emergence in wild birds (Fig. 2a, 2c). Single-segment analyses exhibited greater uncertainty in estimates of B3.13 emergence but remained broadly concordant with the concatenated analysis (Fig. 2c). Since no additional B3.13 viruses outside of the dairy cattle outbreak clade have been observed since early 2024, this lineage appears to not have persisted in the broader wild bird population following its emergence in dairy cattle, with detections in wild birds due to spillbacks from cattle into wild birds and without detected onward transmission. This result highlights how frequently avian influenza reassortants can form, spread, and be subsequently overtaken by later, potentially more fit, reassortant lineages. Further, we find that the A1 genotype from which four segments of B3.13 are derived began to decline around the time that B3.13 was introduced into, and subsequently spread among, dairy cattle (Fig. 2d).

### D1.1 emerged repeatedly in dairy cattle

Approximately one year after the emergence of B3.13 in dairy cattle, the D1.1 genotype was detected in Nevada through surveillance under the National Milk Testing Strategy (Fig. S6). The earliest complete viral genome was sampled from a Nevada dairy herd on 21 January 2025, and a human infection (A/Nevada/10/2025(H5N1)) was identified early in the Nevada outbreak, with the corresponding virus sampled on 14 February 2025. A second D1.1 outbreak was detected in Arizona dairy cattle later that month, with the earliest complete genome sampled on 31 January 2025.

Unlike B3.13, D1.1 was already established in wild birds and had already spilled over repeatedly into poultry and humans before being detected in dairy cattle. We first inferred a maximum likelihood tree of D1.1 genomes from the Arizona and Nevada dairy cattle outbreaks together with a background avian dataset, indicating that these two cattle outbreaks resulted from separate introduction events (Figs. 1, S7).

We next sought to estimate when D1.1 entered the dairy cattle population. Relatively few D1.1 virus genomes have been sampled from cattle (n=43) and humans (n=1), precluding the reliable estimation of an evolutionary rate for this lineage in cattle. We therefore used our estimate of the B3.13 evolutionary rate in dairy cattle and applied this rate to the D1.1 dairy cattle outbreak clades (including the human virus) in a Bayesian phylogenetic framework. The results again confirmed independent introductions of D1.1 into dairy cattle in Nevada and Arizona. For the Nevada outbreak, we inferred that the spillover likely occurred between mid-October 2024 (tPMRCA; 95% HPDI: 15 September 2024–13 November 2024) and early December 2024 (tMRCA; 95% HPDI: 28 November 2024–18 December 2024). For the Arizona outbreak, the corresponding interval spanned late August 2024 (tPMRCA; 95% HPDI: 6 August 2024–28 September 2024) to late December 2024 (tMRCA; 95% HPDI: 24 December 2024–1 January 2025).

The estimated tMRCA of the broader D1.1 lineage was early July 2024 (95% HPDI: 25 June–24 July 2024), consistent with a recent report^13^. As with B3.13, the molecular clock analysis indicates that as little as 1–2 months may have elapsed between the formation of the D1.1 reassortant and spillover into dairy cattle, and that D1.1 subsequently circulated cryptically in dairy cattle for up to 5–6 months prior to its detection. In contrast to B3.13, however, D1.1 has continued to circulate widely among wild birds following its emergence and remains the dominant avian genotype in North America^13^.

### Genome-wide patterns of molecular evolution in dairy cattle

To assess whether the molecular evolutionary dynamics of AIV change following spillover into dairy cattle, we compared evolutionary rates between bovine and avian hosts. We performed codon-aware phylogenetic reconstruction of B3.13 separately in dairy cattle and avian hosts using the A1 concatenated alignment, which improves estimation of overall tree length (Fig. S4), while estimating segment-specific evolutionary rates. We found that the evolutionary rate of B3.13 in dairy cattle was elevated, both in aggregate and for the PA, NA, and MP segments, whereas the evolutionary rate of HA was similar between cattle and avian hosts (Fig. 4a, Table 1).

**Figure 4.**
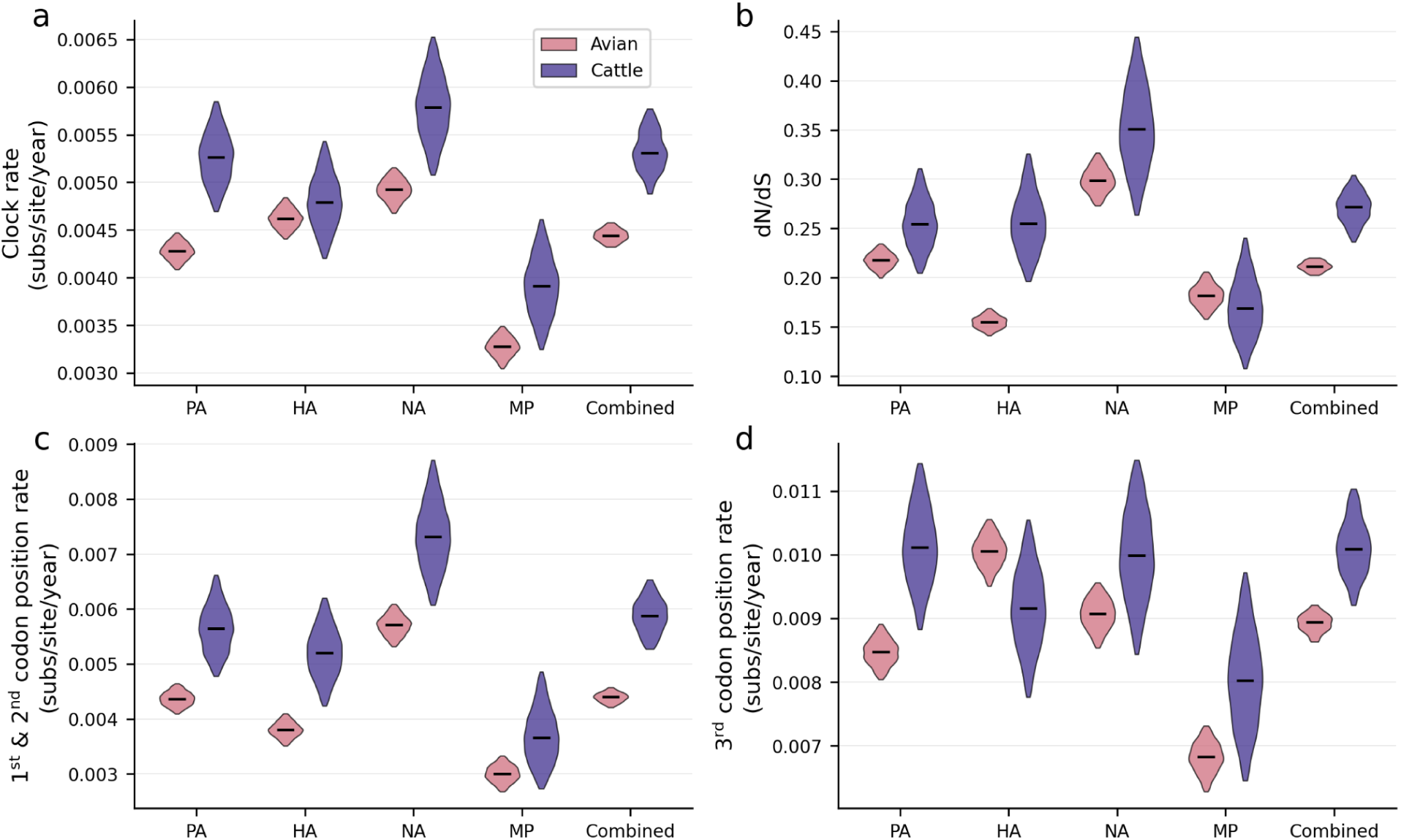
Elevated evolutionary rate and dN/dS of B3.13 in dairy cattle. The evolutionary rate (**a**), dN/dS (**b**), evolutionary rates at the first and second codon positions (**c**), and evolutionary rate at the third codon position (**d**) for the PA, HA, NA, and MP genes, as well as the concatenated four-gene alignment, in avian and cattle hosts. Black lines indicate posterior median values, and violins show the 95% HPDIs. Median and 95% HPDI numerical values are provided in Table S1. “Combined” represents the concatenated PA-HA-NA-MP alignment.

Given the elevated viral evolutionary rate in dairy cattle, we next examined whether selective pressures differed between avian and bovine hosts. We first evaluated genome-wide and segment-specific patterns of selection by estimating the ratio of non-synonymous to synonymous substitution rates (dN/dS), allowing us to assess broad shifts in selective pressure across the viral genome. We then tested whether the intensity of selection differed between hosts and, finally, examined whether host-associated shifts were localized to specific amino acid residues through site-specific analyses.

We estimated dN/dS for B3.13 separately in avian and bovine hosts, both in aggregate and for the four individual segments comprising the concatenated alignment. We found an elevated dN/dS for B3.13 in cattle compared to B3.13 in birds across multiple segments and when averaged across all segments (Fig. 4b, Table 1). The HA segment exhibited the largest difference in dN/dS between hosts, with dairy cattle virus estimates elevated and clearly different from the avian virus estimates (*i.e.*, non-overlapping 95% HPDIs). The dN/dS values of PA and NA were also elevated in cattle, whereas the dN/dS of MP was similar to, or slightly lower than, that observed in avian hosts. The elevated dN/dS observed in dairy cattle was driven by increased substitution rates at the first and second codon positions relative to the third codon position, across both the concatenated alignment and its constituent segments (Figs. 4c–d, S8, Table 1).

We next assessed whether D1.1 exhibited a similar host-associated shift in dN/dS. Given the limited number and diversity of D1.1 genomes from dairy cattle, we estimated dN/dS using a single codon-aware phylogenetic reconstruction and compared values between avian and bovine lineages based on inferred counts of non-synonymous and synonymous substitutions and their neutral expectations. Specifically, we performed within-tree comparisons of dN/dS between avian branches and two cattle-associated branch sets: the dairy cattle outbreak clades and their stem branches. For the dairy cattle outbreak clades, the average fold change in dN/dS relative to the avian background exceeded one (Fig. 5), indicating a directional shift following host switching. Support for a fold change greater than one was modest when considering only the dairy cattle outbreak clades (Fig. 5a; posterior probability = 0.77) but increased when stem branches were included (Fig. 5b; posterior probability = 0.91). Similarly, the stem branches showed a directional increase relative to the avian background (Fig. 5c; posterior probability = 0.82).

**Figure 5.**
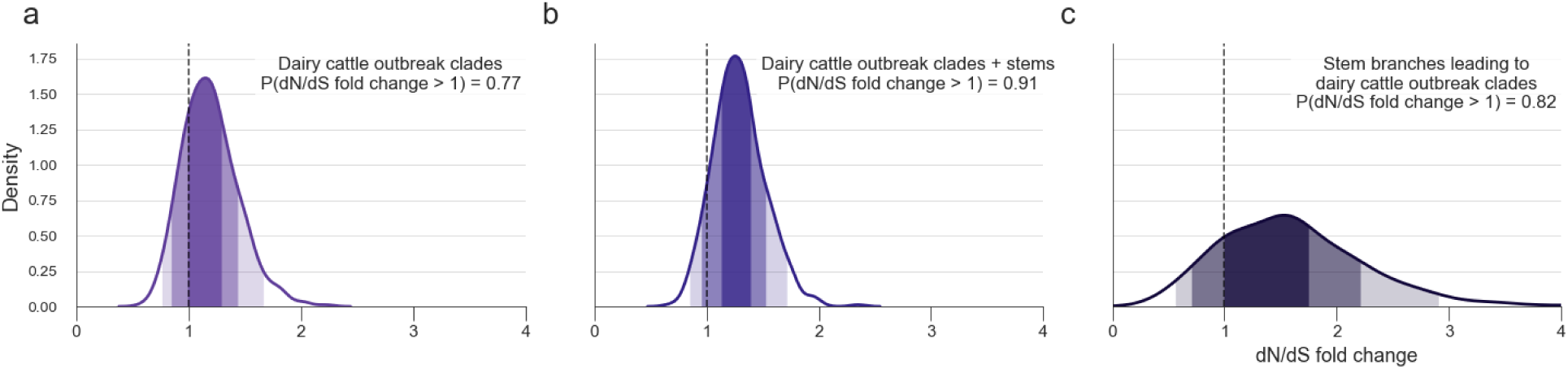
Elevated dN/dS of D1.1 in dairy cattle. Fold change in dN/dS relative to the avian background for the dairy cattle outbreak clades (**a**), for the outbreak clades together with their stem branches (**b**), and for the stem branches alone (**c**). For all panels, lightest shading indicates the 95% HPDI, medium shading the 80% HPDI, and darkest shading the 50% HPDI. Dashed lines at 1.0 in panels indicate no fold change and we report the posterior probability P(dN / dS fold change > 1) relative to a prior probability of 0.5.

When examining dN/dS estimates across the posterior distribution of trees (as opposed to within-tree comparisons), we found that, although the posterior distributions overlapped substantially, the mean and median dN/dS estimates for the dairy cattle outbreak clades were higher than those for the avian background, with HPDIs shifted upward (Fig. S9). Including the stem branches together with the dairy cattle outbreak clades modestly increased dN/dS estimates, yielding a non-overlapping 50% HPDI relative to avian hosts. The average dN/dS values for the stem branches exceeded those of the avian background (Fig. S9), although the associated HPDIs were wide, reflecting substantial uncertainty given that these estimates are based on only two branches. Altogether, these analyses indicate that D1.1 exhibits a host-associated shift in selective pressure consistent with that observed for B3.13 and suggest that the stem branches may represent, at least in part, viral evolution in dairy cattle.

### Evidence for relaxed purifying selection in dairy cattle

Following spillover from avian into bovine hosts, H5N1 exhibited evidence of an altered selective environment. However, shifts in dN/dS can arise from multiple evolutionary processes, including changes in selection intensity or shifts in the balance between purifying and diversifying selection. The elevated dN/dS values observed for B3.13 and, likely, D1.1 could reflect an increase in diversifying selection, a reduction in the strength of purifying selection, or a combination of the two.

To distinguish between these possibilities, we applied the RELAX modeling framework^14^ to test whether viruses circulating in dairy cattle experienced changes in selection relative to avian viruses, and to assess whether any such shifts were driven by changes in purifying or diversifying selection. We modelled selection regimes for two phases of the B3.13 and D1.1 outbreaks, partitioning the phylogenetic trees into three sets of branches: (i) branches representing evolution in the wild bird reservoir, (ii) branches representing evolution during the dairy cattle outbreaks, and (iii) the stem branches separating these host-associated clades, including potential evolutionary intermediate lineages (Fig. S10; see Methods). Briefly, we estimate a single scaling parameter *K* that modifies the background distribution of dN/dS values: *K* < 1 indicates a shift toward neutrality, consistent with relaxed selection in dairy cattle, whereas *K* > 1 indicates intensification of selection away from neutrality, strengthening both purifying and diversifying selection.

We examined selection dynamics in the A1 sublineage, corresponding to HA, PA, NA, and MP segments, that led to the B3.13 reassortant, by comparing the avian background with the dairy cattle outbreak clade (Fig. S10a). We found strong support for a relaxation of selection on branches corresponding to the dairy cattle outbreak relative to the avian background (*K* = 0.40; *p* < 0.0001; Fig. 6b). Consistent with our Bayesian phylodynamic inference (Fig. 4), the inferred aggregate dN/dS during the first year of circulation in dairy cattle exceeded that observed in avian hosts (Fig. 6a). This elevation in dN/dS was likely driven by a relaxation of purifying selection rather than an intensification of diversifying selection (Fig. 6c), indicating that H5N1 is evolving in distinct selective regimes in avian versus bovine hosts. We also observed evidence for relaxed selection in PB2, NP, and NS, although PB1 did not show a significant change in selection intensity between avian and bovine hosts (Fig. 6b–c, Table S2).

**Figure 6.**
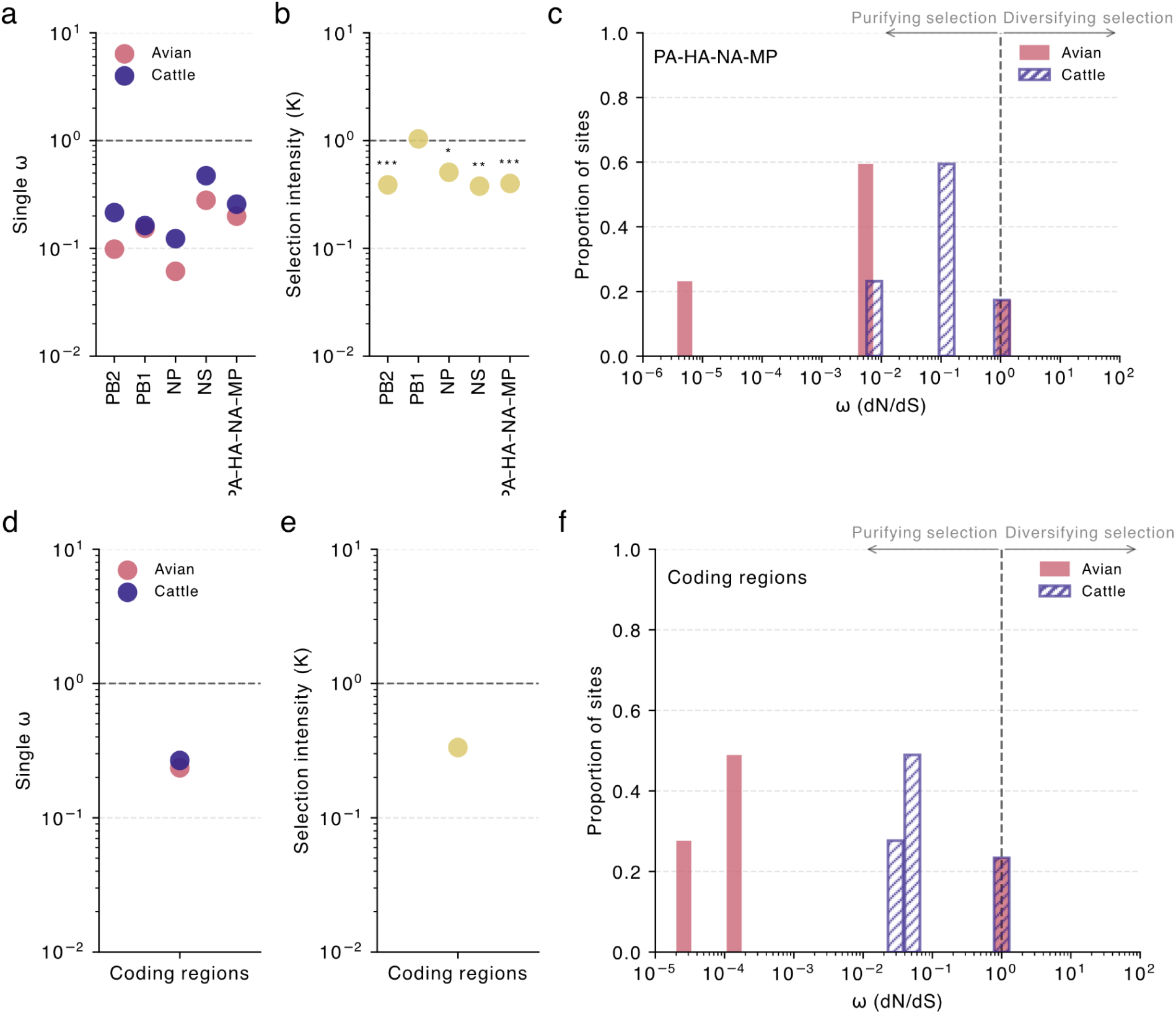
Relaxed selection of H5N1 in dairy cattle. **a**, Single ω point-estimates for the avian and cattle partitions of the branch sets for the concatenated alignment and the four remaining segments. For each segment, estimates are based on the segment-specific phylogeny reflecting its reassortment history, with all lineages corresponding to the B3.13 dairy cattle outbreak at the whole-genome level (see Figure S1). **b**, Change in selection intensity between avian and cattle branch sets for the concatenated alignment and the four remaining segments, based on the same segment-specific phylogenies. **c**, Three ω categories and their corresponding proportions of sites for avian and cattle branch sets for the concatenated alignment. Each bar corresponds to an ω category and proportion of sites. **d**, Single ω estimates for the avian and cattle partitions of the branch sets for D1.1. **e**, Change in selection intensity between avian and cattle branch sets for D1.1. **f**, Three ω categories and their corresponding proportions of sites for avian and cattle branch sets for D1.1. Numerical values for each of the four individual segments and the concatenated alignment relevant to B3.13, as well as for D1.1, are provided in Table S2. Statistical significance is indicated by * p < 0.05, ** p < 0.01, and *** p < 0.001. Refer to Figure S10 for schematics indicating how B3.13 and D1.1 branches were partitioned.

With the D1.1 lineage, we compared the avian background with the two dairy cattle outbreak clades, which we analyzed as a combined group (Fig. S10b). We observed a trend consistent with relaxed selection in dairy cattle (*K* = 0.33; Fig. 6e); however, this result was not strongly supported (*p* = 0.5629), likely reflecting the limited number of available D1.1 cattle virus genomes. Despite this limited power, the inferred shift in selection appeared driven by a reduction in purifying selection rather than an increase in diversifying selection (Fig. 6f). This pattern is consistent with the shift in selection regime observed for B3.13 following spillover into dairy cattle.

### Site-specific signatures of selection

The genome-wide and gene-specific analyses indicate that H5N1 experienced altered selective pressures following its introduction into dairy cattle. To determine whether specific sites were subject to differential selective pressures in avian versus bovine hosts, we performed selection analyses at individual amino acid sites across all eight gene segments in both hosts. We focused on the B3.13 outbreak, for which greater sampling in dairy cattle provides increased statistical power. Site-specific patterns were evaluated across the entire viral genome (Tables S3–S6; Supplemental Note 1), and here we highlight the HA segment, which exhibited the clearest host-associated differences.

Using the robust counting framework^15^ in BEAST X^16^, we identified 14 sites under positive selection and 504 sites under negative selection in HA in dairy cattle (Fig. 7a; Table S3), while 16 sites were inferred to be under positive selection and 517 sites under negative selection in avian hosts (Fig. 7b; Table S3). In bovine viruses, sites under positive selection were enriched in the HA1 domain, in particular the globular head domain, whereas in avian hosts, positively selected sites were more frequently located flanking this region or near the 5’ and 3’ ends of the gene. The majority of sites in both hosts were inferred to be either neutrally evolving or under purifying selection. Site-specific selection patterns inferred using the FUBAR framework^17^ corroborated our results in dairy cattle but identified fewer sites under positive selection in avian hosts than did the robust counting framework (Fig. 7, S11; Table S4). Sites inferred to be under positive selection in cattle but neutral or negatively selected in birds exhibited substantially higher dN/dS values in dairy cattle, and an analogous pattern was observed for sites under positive selection in birds but not in cattle. Sites inferred to be under positive selection in both bovine and avian hosts exhibited moderately elevated but broadly similar dN/dS values across hosts (Fig. 7c). Although the HA segment has the most positively selected sites in avian and bovine hosts combined, these patterns are broadly consistent among the other segments (Tables S3, S4).

**Figure 7.**
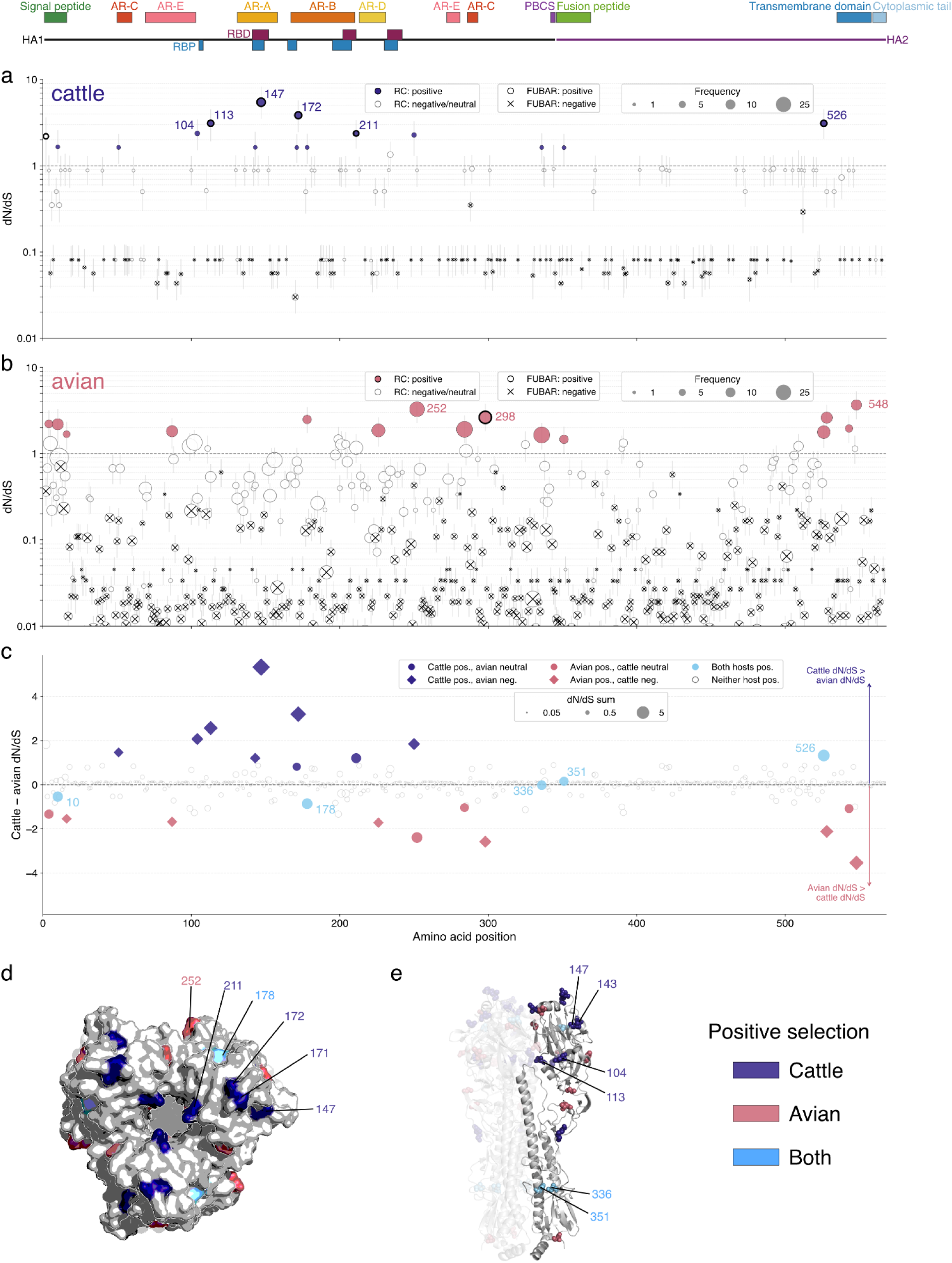
Site-level selection for HA in the A1 lineage in avian and dairy cattle hosts. Site-specific dN/dS estimates for the HA gene for the A1 lineage in (**a**) dairy cattle and (**b**) avian hosts, based on the robust counting framework. The A1 subclade circulating in dairy cattle corresponds to the B3.13 outbreak at the whole-genome level. Scatter points indicate posterior medians and are sized by the average frequency of mutation at each site in the HA phylogeny; light grey lines indicate the 95% HPDI. Sites with 95% HPDIs entirely above 1 are colored red in dairy cattle and blue in avian hosts. A thick black outline indicates positive selection inferred by FUBAR at that site in the corresponding host, while a cross indicates negative selection inferred by FUBAR. Highly selected sites are annotated. **c**, Difference between median dairy cattle and avian dN/dS values per site, with points sized by the sum of the median dairy cattle and avian dN/dS values. Point shape and color indicate whether one, both, or neither host shows positive, neutral, or negative selection. Sites that are neutral or negatively selected in both hosts are shown as light grey circles. Sites under positive selection in both hosts are annotated. **d**, Surface representation of H5 from the human H5N1 strain A/Texas/37/2024 (PDB: 9DWE) viewed from the top, with positively selected sites colored by host. **e**, Side view of the H5 structure, with positively selected sites shown as spheres and colored by host. HA domains are indicated in the schematic at the top. Antigenic regions (A–E) are denoted as AR. PBCS, polybasic cleavage site; RBD, receptor-binding domain; RBP, receptor-binding pocket. HA sites are numbered using the sequential numbering scheme; see Table S5 for conversion.

Sites inferred to be under positive selection in both bovine and avian hosts exhibited higher substitution frequencies over time compared with other sites, with this pattern being more pronounced among sites under the strongest positive selection in dairy cattle. Many of these sites experienced repeated substitutions across the phylogeny, and two of the sites inferred to be most strongly selected in dairy cattle—positions 104 and 147 (HA sites are numbered using the sequential numbering scheme; see Table S5 for conversion)— have been associated with antigenic escape^18,19^. Position 147, in particular, exhibited a disproportionately high number of substitutions over time. More broadly, and after adjusting for the total number of substitutions observed in each host, sites inferred to be under positive selection in dairy cattle accumulated substitutions more frequently than did sites under positive selection in wild birds (Fig. S12).

Additionally, although the number of sites under positive and negative selection does not appear to have substantively shifted following the emergence of H5N1 in dairy cattle, the strength of purifying selection among negatively selected sites differs markedly between hosts (Fig. 7a–b). Sites classified as negatively selected in cattle have dN/dS values closer to 1 than do negatively selected sites in birds (Fig. S13). The degree of positive selection at positively selected sites, by contrast, appears broadly similar across hosts (Figs. 7a–b, S13). This pattern is consistent with a relaxation of purifying selection after host-switching, corroborating our RELAX results (Fig. 6).

Site-specific analyses of the remaining gene segments also revealed host-associated differences in selection patterns (Tables S3–S6; Figs. S15–S18; Supplemental Note 1). In several cases, sites exhibited opposing selection regimes between hosts, appearing under positive selection in one host but negative selection in the other (Fig. S14; Supplemental Note 1). One notable site inferred to be under positive selection in bovine hosts but under negative selection in avian hosts is PB2 position 740, where D740N has been shown to boost polymerase activity in cattle cells in a cattle H5N1 background^20,21^.

## Discussion

In reconstructing the emergence and evolution of B3.13 and D1.1 in dairy cattle and wild birds, we find that each outbreak occurred within several months of the appearance of these reassortant genotypes in avian hosts, with cryptic circulation of each lineage in dairy cattle before detection. Our results suggest that these lineages evolve more rapidly and with fewer selective constraints in dairy cattle than in birds, and that continued circulation in dairy cattle has resulted in repeated amino acid substitutions at sites previously implicated in mammalian adaptation.

By performing multi-segment analyses and accounting for host differences in our modeling, we obtained a narrower—and more recent—estimate of B3.13 spillover into dairy cattle compared to previous HA-focused analyses^11,22^. We also estimated elevated evolutionary rates in dairy cattle relative to birds that fall within the expected range for influenza A viruses, whereas previous estimates^11,22^ were nearly an order of magnitude higher. This difference likely reflects improved phylogenetic resolution when using concatenated multi-segment analyses. Together, these results highlight the importance of multi-segment analyses when reconstructing the emergence and evolution of reassorting viruses.

The emergence of HPAIV in US dairy cattle has reshaped our understanding of AIV ecology, transmission, and host range. The detection of B3.13 and D1.1 in cattle within months of their emergence in avian hosts suggests that spillover from wild birds into cattle may not be rare, but that only a subset of such introductions lead to sustained transmission. For example, a third independent spillover of D1.1 into dairy cattle was detected in Wisconsin in December 2025 (ref.^23^), but it does not appear to have resulted in onward spread beyond a single premise. Sustained transmission in the US may be facilitated by the highly interconnected nature of the dairy cattle network^24,25^, even though the precise mechanisms of transmission remain incompletely understood. If H5N1 spillovers from wild birds into cattle also occur in other countries^26^, lower levels of cattle connectivity elsewhere could help explain why sustained transmission has not been widely observed outside the US. Collectively, the four detected spillovers of H5N1 into US dairy cattle illustrate a range of outcomes, from limited transmission within a single herd to sustained multi-regional spread, likely shaped by differences in network structure, surveillance intensity, and intervention measures such as mandated interstate testing.

The B3.13 lineage showed limited spread among wild birds but continued transmission among dairy cattle, whereas D1.1 exhibited limited spread in dairy cattle but sustained transmission among wild birds. This contrast indicates that fitness differences of viral lineages observed in one host do not necessarily translate across hosts. Most of the key substitutions associated with mammalian adaptation arose early during the initial expansion of the B3.13 outbreak in northern Texas (e.g., PB2:M631L and PA:K497R; refs.^21,27^). By the time the virus spread to other states, it already appeared capable of sustained transmission in dairy cattle, with little evidence for selection of additional canonical PB2 mammalian adaptations (e.g., D701N, E627K, Q591K; refs.^21,28^). In contrast, the D1.1 lineage experienced sustained transmission among dairy cattle in Arizona for more than 6 months, despite the absence of canonical mammalian adaptations (the D1.1 virus that infected Nevada cattle had the well-described D701N adaptation^20^). In the genome sequences of influenza A virus spillover into swine where onward transmission has been established (the Eurasian-avian swine lineage), and in pinnipeds where it has been inferred (the 2.3.4.4b B3.2 lineage in South America), the spectrum of mutations seen largely involve different amino acid residues^29,30^. The only shared cattle-associated substitution in these species that we found is PB2:D701N, but when considering the broader spillover events of multiple H5 and LPAI viruses into pinnipeds, the changes involve overlapping residues with the cattle lineages but do not always result in the same amino acid (*e.g.*, PB1:S384P, PA:I448V/I and E613K/T; NS1:S7L/T and potentially L77R/F [ambiguous codon in pinniped sequence]). There may be more evolutionary pathways to transmission fitness in mammals in general, and cattle specifically, than have been documented thus far. These observations suggest that viral fitness following spillover is host specific and may depend on broader genomic and epidemiological context rather than a fixed set of canonical mammalian adaptations.

Our results suggest that, upon entering dairy cattle, H5N1 has experienced a reduction in the intensity of purifying selection and is evolving more rapidly than in wild birds, a pattern that has been observed in the early stages of epidemics and pandemics^31^. The virus is therefore better able to explore fitness landscapes and sample mutations that may confer increased fitness. Despite two years of circulation in cattle and potential reinfection of the same herds and animals, it remains unclear to what extent the transmission potential of B3.13 has shifted over time. There is limited evidence for evolution at HA epitopes that are important for antigenic change in other host species (e.g., humans and swine), although two substitutions—D104G and V147M—have been described as antigenically relevant in recent studies^18,19^. Further, sites inferred to be under positive selection in HA in dairy cattle are concentrated in regions proximal to the receptor binding site, potentially driven by immunity or receptor binding. Previous spillovers of avian influenza viruses into new hosts have shown that adaptive evolution can continue for many years after introduction; for example, the Eurasian lineage circulating in swine accumulated adaptive substitutions in HA and NA for a decade after its emergence^32^, and our results here indicate that diversifying selection has not substantively shifted after H5N1 emerged in dairy cattle. Continued circulation of B3.13 in California and other states could increase immune pressure on the virus over time. Antigenic evolution will be important to monitor, particularly if an H5N1 vaccine is licensed for use in US dairy cattle, as both the rate of evolution and the selective environment could shift^33^.

The detail with which we document the evolution of H5N1 in dairy cattle and wild birds provides an opportunity to re-examine how evolutionary dynamics change following sustained establishment in a new host. Our results demonstrate that relaxed selective constraints, host-specific fitness landscapes, and genomic context jointly shape viral evolution after host shifts. Together, these analyses establish a baseline for interpreting ongoing and future evolution of H5N1 in dairy cattle. Continued monitoring of the genomic evolution of H5N1 in dairy cattle will therefore be essential for determining how these evolutionary patterns change over time, particularly as immune pressure and control measures evolve.

---

**Figure S1.**
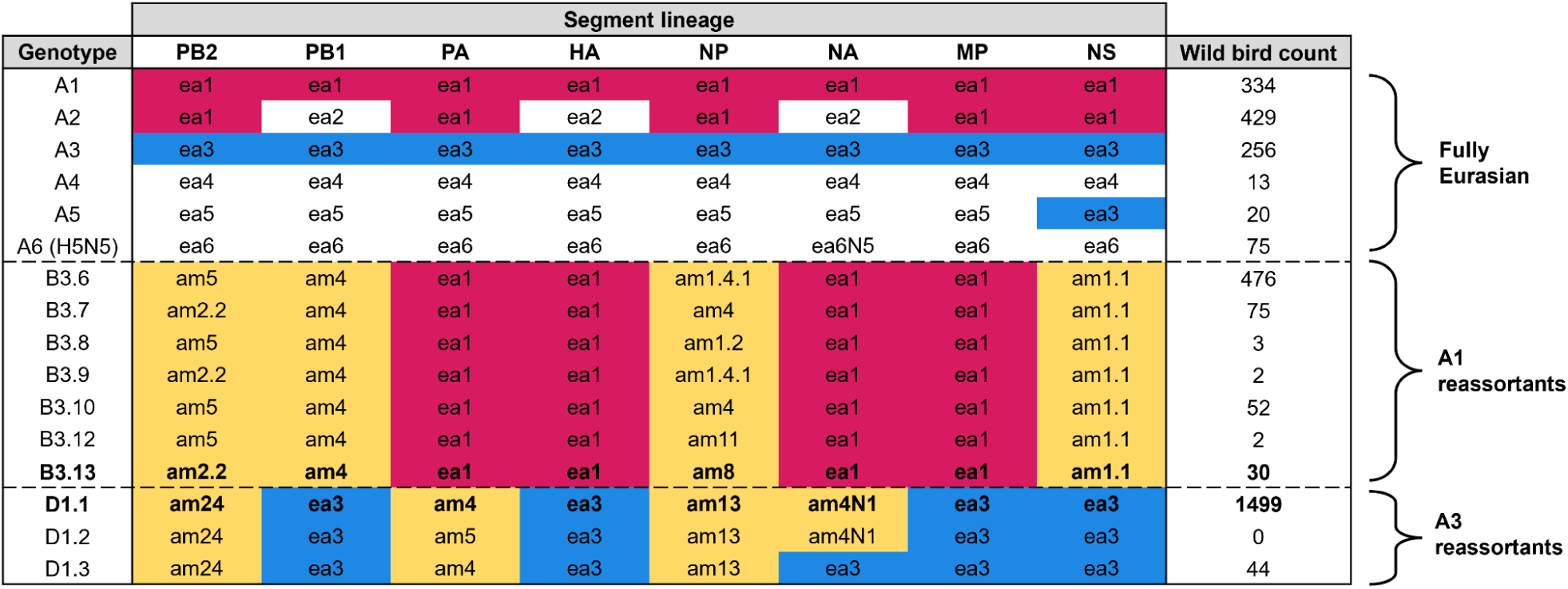
Gene segment lineage constellations defined by GenoFLU for Eurasian clade 2.3.4.4b genotypes (A1-A6), North American A1 reassortant genotype B3.6 and its descendants (including B3.13), and North American A3 reassortant genotypes D1.1, D1.2 and D1.3. The geographical origin of each lineage is denoted as either ‘ea’ (Eurasian) or ‘am’ (American). The ancestral Eurasian segments for the ‘B’ and ‘D’ genotypes are shaded in pink and blue, respectively, while segments of North American low pathogenicity AIV (LPAIV) origin are shaded in yellow. Wild bird sample counts for each genotype are as of June 13, 2025. Genotype D1.2 has thus far only been found in poultry and swine.

**Figure S2.**
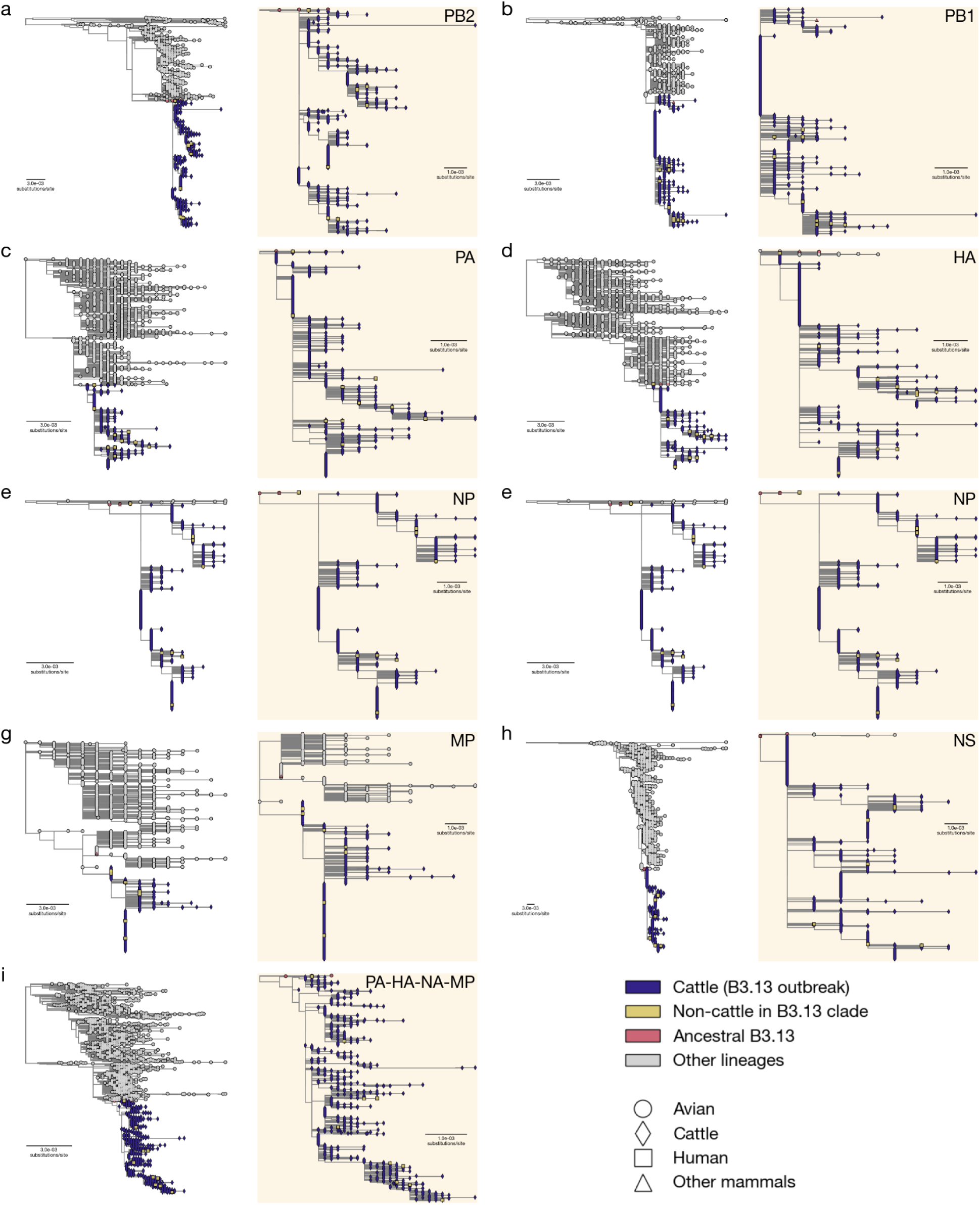
Maximum likelihood trees for each H5N1 gene segment (a–h) and the concatenated alignment composed of PA, HA, NA, and MP (i). The dairy cattle outbreak clade is highlighted in the yellow box on the right of each panel. Lineage is indicated by scatter point color and host is indicated by scatter point shape, with human virus genomes represented as yellow squares. The earliest human B3.13 sequence (A/Texas/37/2024(H5N1)) was sampled from a dairy worker on 28 March 2024, 21 days after the earliest sampling of a poultry case (7 March 2024) and 18 days after the earliest dairy cattle case was sampled (10 March 2024). This human virus genome is not consistently nested with the dairy cattle outbreak clade, clustering with cattle viruses only in the NA, PB1, NS, and MP segments. In MP, the human virus genome is genetically identical to many early cattle virus genomes. In the remaining segments, the human genome contains, in aggregate, 8 additional derived substitutions relative to the basal cattle lineages, including PB2 E627K, a canonical mammalian adaptation. Of these, 7 are unique, and the remaining substitution (NP: C936T) is shared with the bovine viruses. However, this homoplasic substitution is likely due to phylogenetic uncertainty, as a whole genome maximum likelihood reconstruction of B3.13 results in the human virus clustering with the dairy cattle outbreak clade rather than with an avian virus (see Figure S3). Collectively, these results suggest that the human virus is not ancestral to the dairy cattle outbreak but sibling to it.

**Figure S3.**
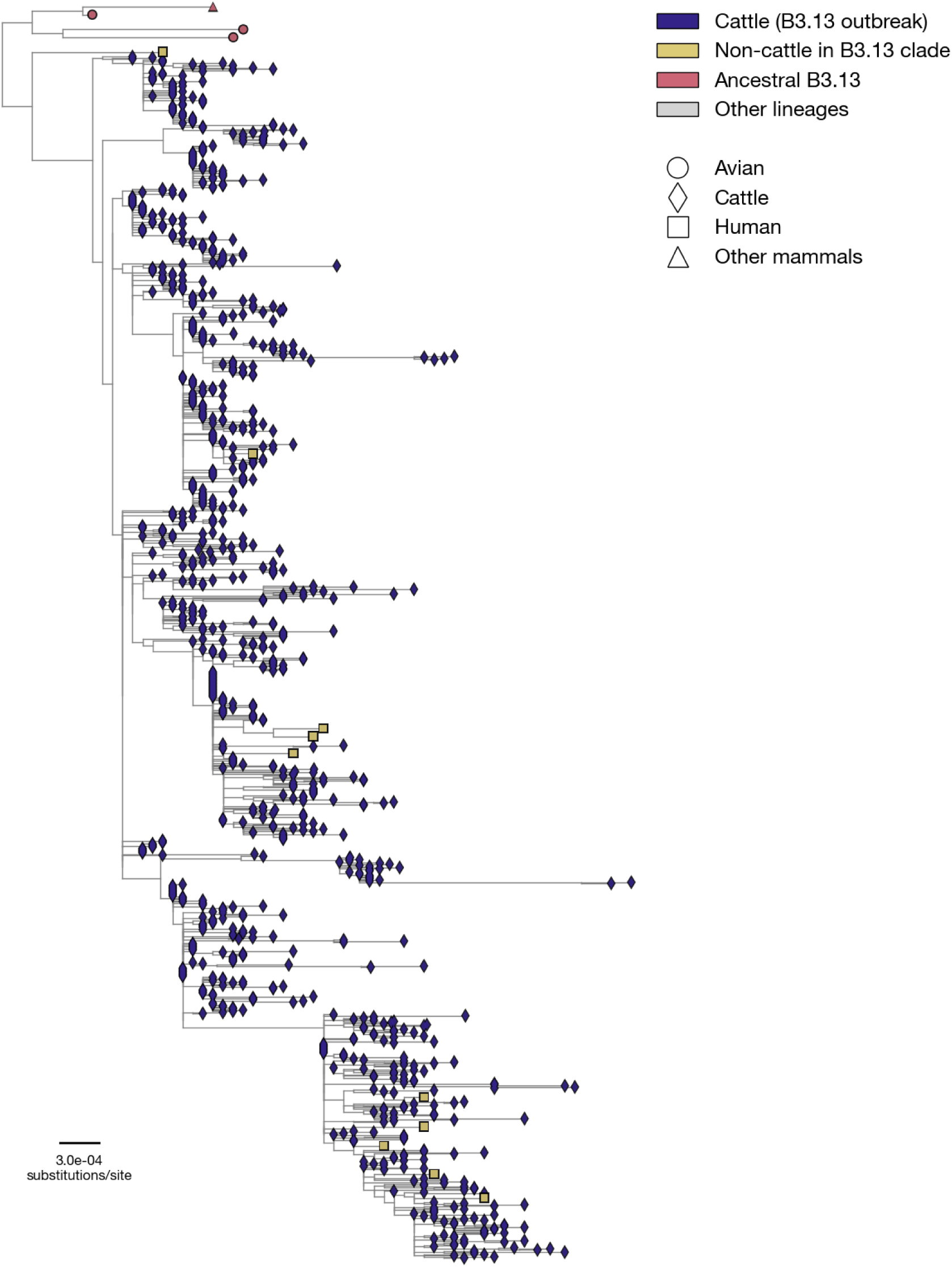
Maximum likelihood tree for a concatenated B3.13 alignment composed of each segment. Lineage is indicated by scatter point color and host is indicated by scatter point shape. Only four B3.13 genomes appear distinct from the dairy cattle outbreak, sampled from a goose in Colorado, a Canada goose in Wyoming, a peregrine falcon in California, and a skunk in New Mexico; each of these was sampled between late November 2023 and late February 2024, before detection of B3.13 in either cattle or poultry. Their phylogenetic placements indicate that these wildlife viruses represent sibling, rather than ancestral, lineages of the dairy cattle outbreak.

**Figure S4.**
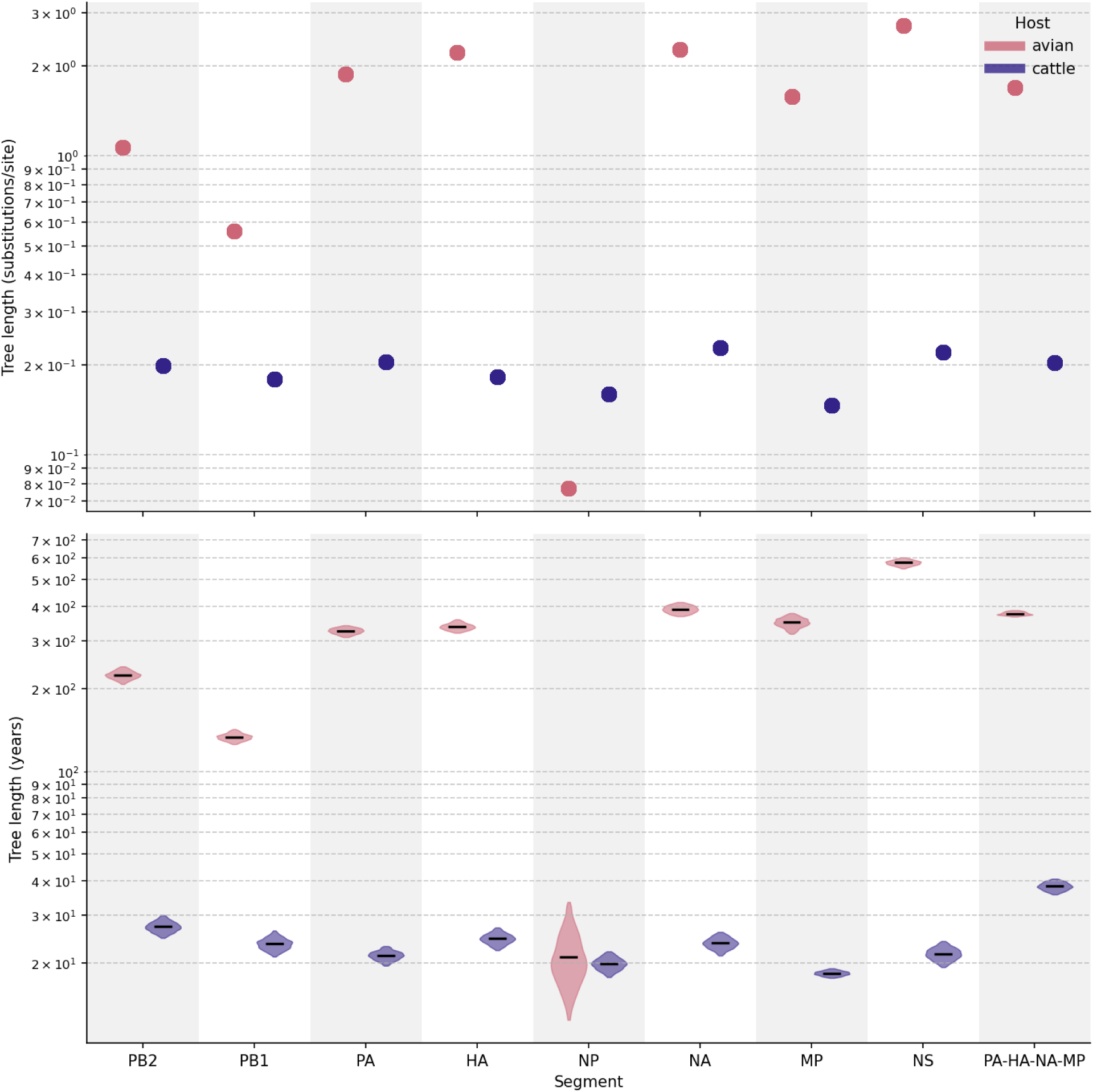
Tree length from mutation (top) and time-calibrated (bottom) phylogenetic trees, for each segment and the A1 concatenated alignment composed of PA, HA, NA, and MP. The black line represents the posterior median time-calibrated tree length, with violins indicating the 95% HPDIs. Among phylogenies inferred from the dairy cattle outbreak virus genomes, maximum likelihood tree lengths were similar across analyses; however, the time-calibrated phylogeny inferred from the concatenated alignment exhibited a greater tree length than those inferred from individual segments. In contrast, for phylogenies inferred from the avian background dataset, both maximum likelihood and time-calibrated tree lengths followed similar relative patterns, likely reflecting the greater temporal and phylogenetic signal present in these more diverse datasets. Together, these results underscore the importance of using concatenated alignments that capture identical evolutionary histories when possible, rather than relying on single segments alone—particularly when inferring early outbreak dynamics from datasets with limited genetic diversity.

**Figure S5.**
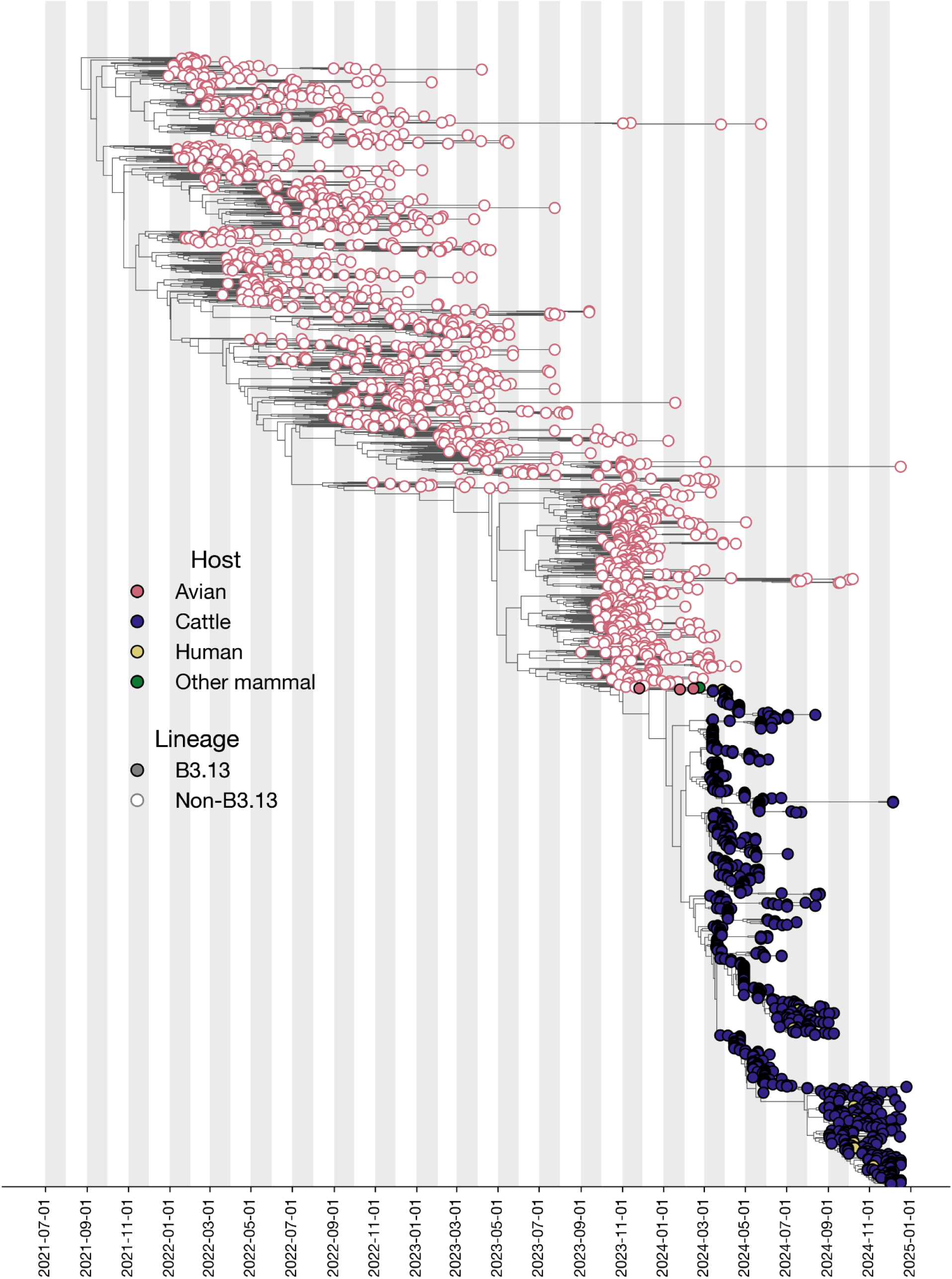
Time-calibrated phylogeny of the A1 lineage inferred from a concatenated alignment of the PA, HA, NA, and MP segments. The phylogeny includes the B3.13 dairy cattle and human viruses and background avian influenza viruses from the broader A1 lineage (see Methods). Although B3.13 is defined at the whole-genome level, the four concatenated segments fall within the A1 lineage. Tips are colored by host, with lineage membership indicated by filled versus open points.

**Figure S6.**
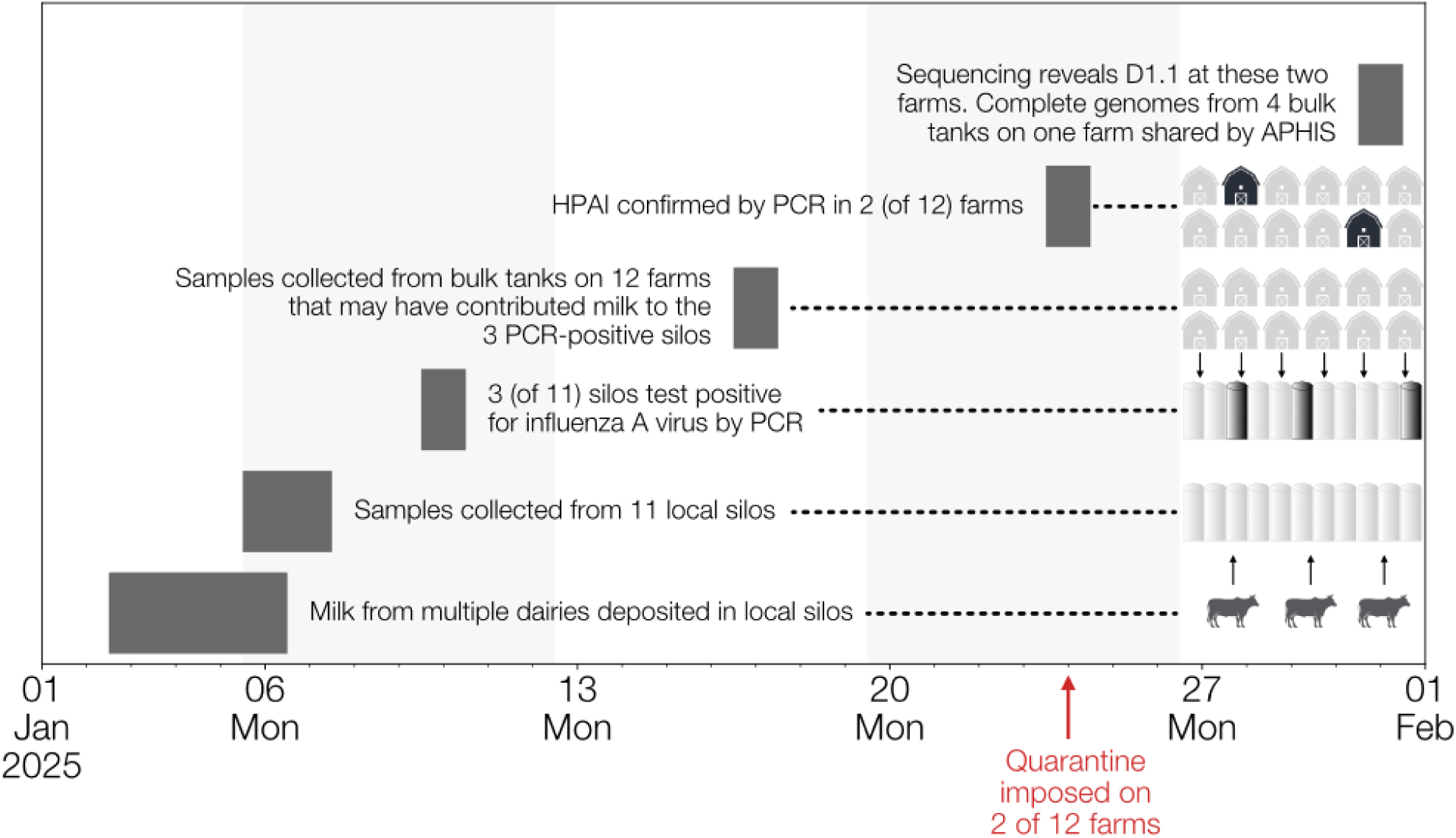
Timeline of D1.1 emergence in cattle in Northern Nevada. Timeline of surveillance and diagnostic events associated with detection of H5N1 clade 2.3.4.4b genotype D1.1 in Churchill County, Nevada. In early January 2025, highly pathogenic avian influenza (HPAI) virus was detected in bulk milk samples collected from dairy processing plant silos under the National Silo Monitoring Program, part of the US Department of Agriculture’s National Milk Testing Strategy^34^. Samples from three of eleven silos collected on 6–7 January 2025 tested positive by PCR at the National Veterinary Services Laboratories (NVSL) on 10 January. Because milk may be stored in silos for up to 72 hours^35^, these samples could have included milk deposited as early as 3 January 2025. Trace-back investigations identified 12 dairies that may have contributed milk to the positive silos. On-farm bulk milk samples collected on 17 January were confirmed HPAI-positive in two dairies on 24 January. Whole-genome sequencing completed on 31 January identified H5N1 clade 2.3.4.4b genotype D1.1 in four bulk tanks from one herd, with a partial D1.1 sequence recovered from a second herd. Large wild bird die-offs were reported near the affected dairies, although the timing of these events and whether avian samples were tested are unknown.

**Figure S7.**
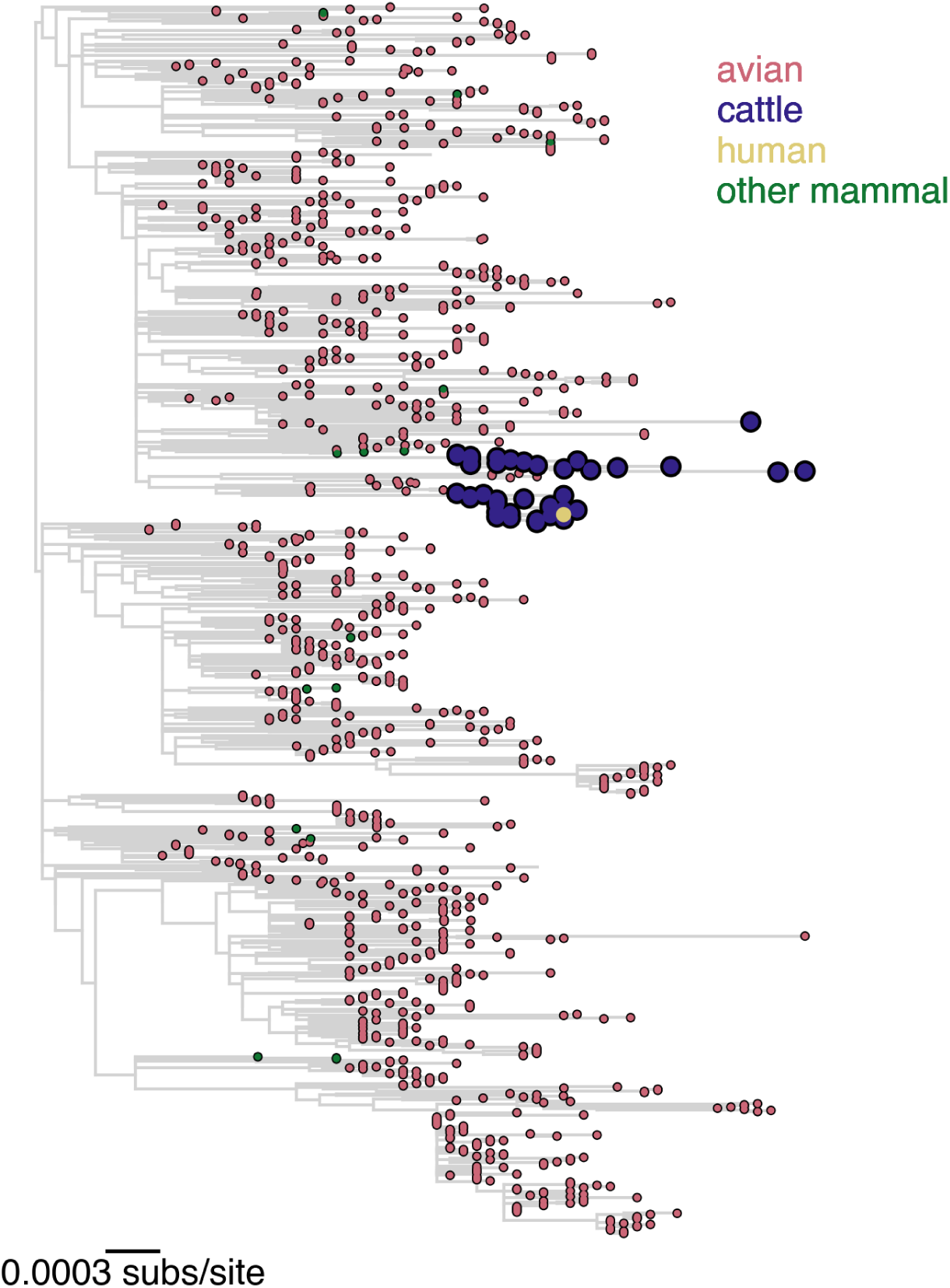
Maximum likelihood phylogeny of the 2.3.4.4b D1.1 lineage genomes. Tips are colored by host: avian (red), cattle (purple), human (yellow), other mammal (green).

**Figure S8.**
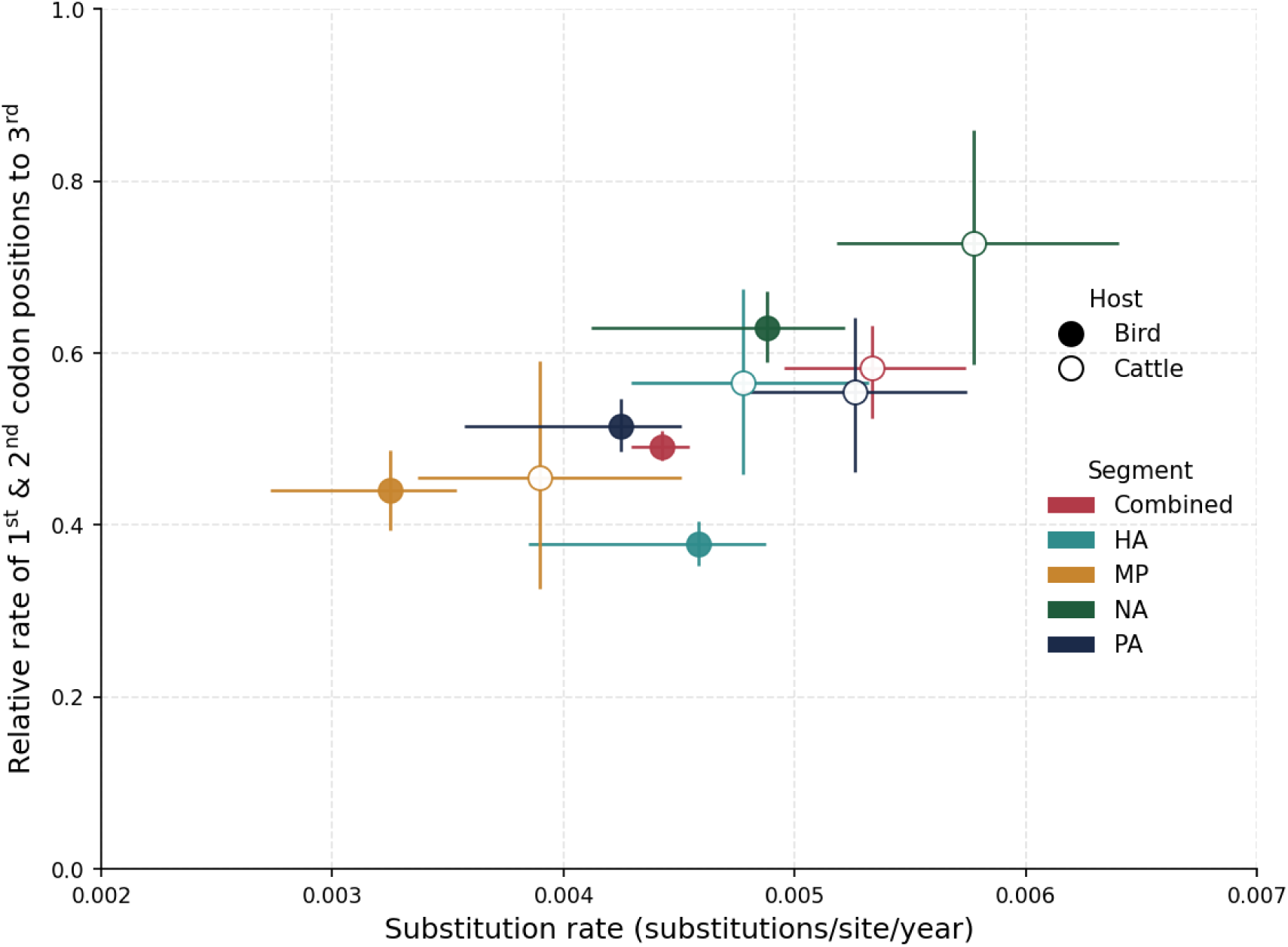
Rates of nucleotide substitution for four genome segments of the A1 lineage and for their concatenated alignment, shown for both avian and dairy cattle hosts. In dairy cattle, the A1 subclade corresponds to the B3.13 outbreak at the whole-genome level. Evolutionary rates are measured as the total number of nucleotide substitutions per site per year (x-axis) and the relative substitution rate at the first and second codon positions compared with the third codon position (y-axis).

**Figure S9.**
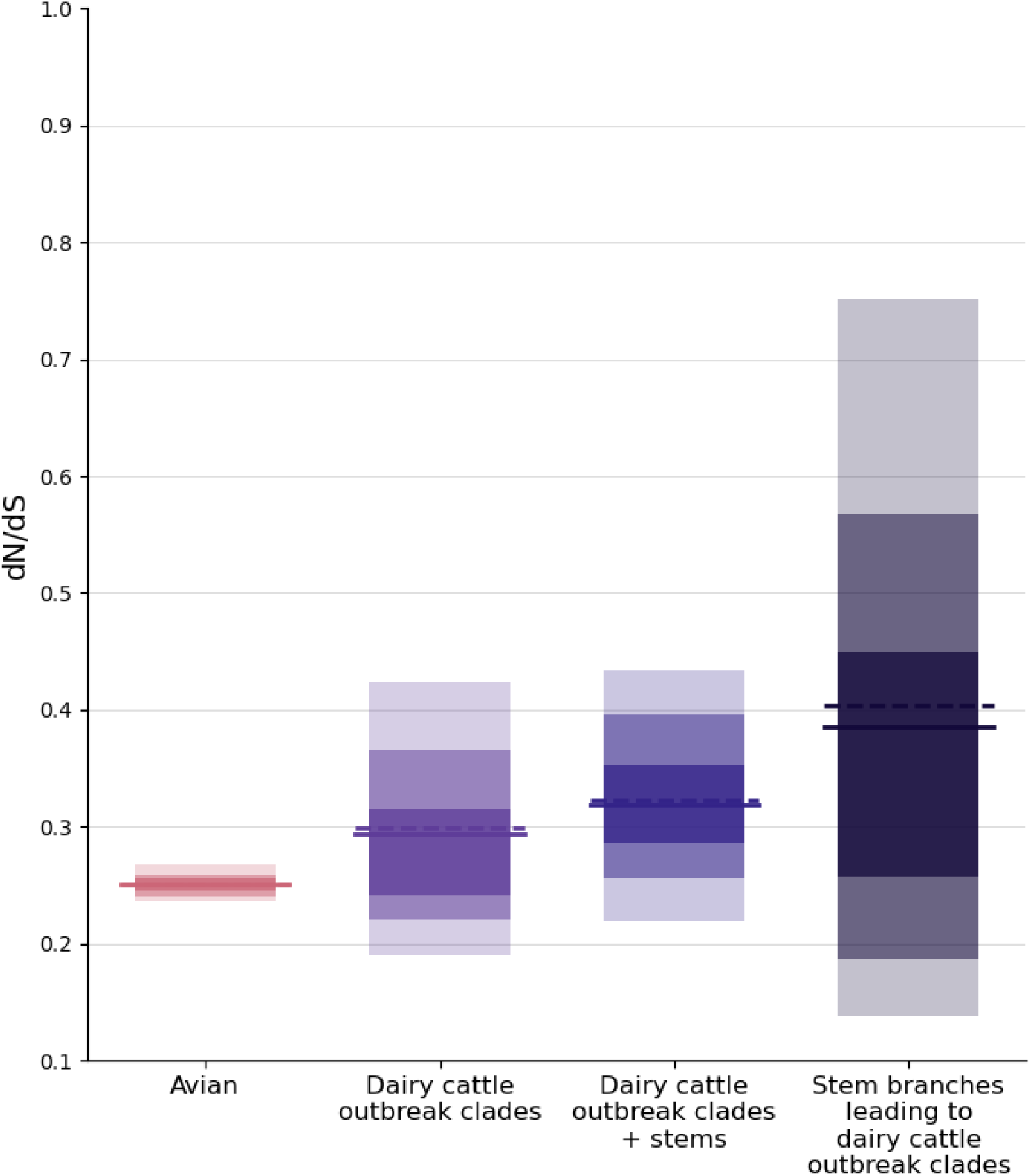
Posterior density of dN/dS for the avian background of the D1.1 phylogeny, the two dairy cattle outbreaks combined, the two dairy cattle outbreaks together with their stem branches, and the stem branches alone. Lightest shading indicates the 95% HPDI, medium shading the 80% HPDI, and darkest shading the 50% HPDI. Solid line represents median dN/dS; dashed line represents mean dN/dS. The mean and median dN/dS values are the same for the avian background; the mean dN/dS is higher for each other dataset.

**Figure S10.**
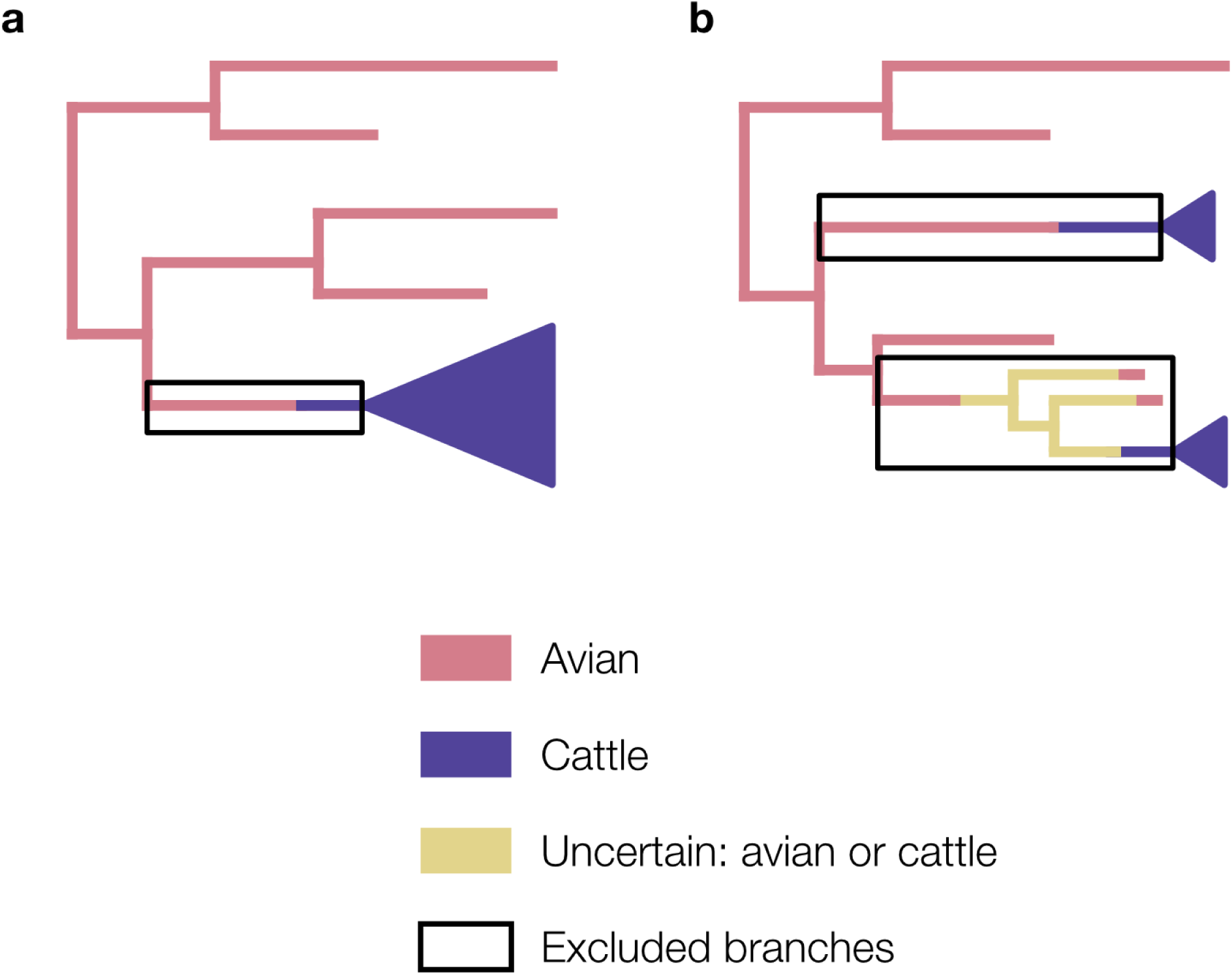
Schematics of the (a) B3.13 and (b) D1.1 phylogenies illustrating host switching. For B3.13, a single branch in the phylogeny plausibly represents the host switch into dairy cattle. For D1.1, there are multiple branches that could reflect evolution in either the avian reservoir or within the dairy cattle outbreak, as several virus genomes cluster with the cattle outbreak and are sampled from domesticated animals or poultry. These genomes may represent independent introductions from wild birds or cryptic transmission associated with the cattle outbreak. Branches corresponding to host switching from wild birds to dairy cattle, or to uncertain avian or bovine host assignment, are excluded from the avian and dairy cattle outbreak datasets and are indicated by black outlines.

**Figure S11.**
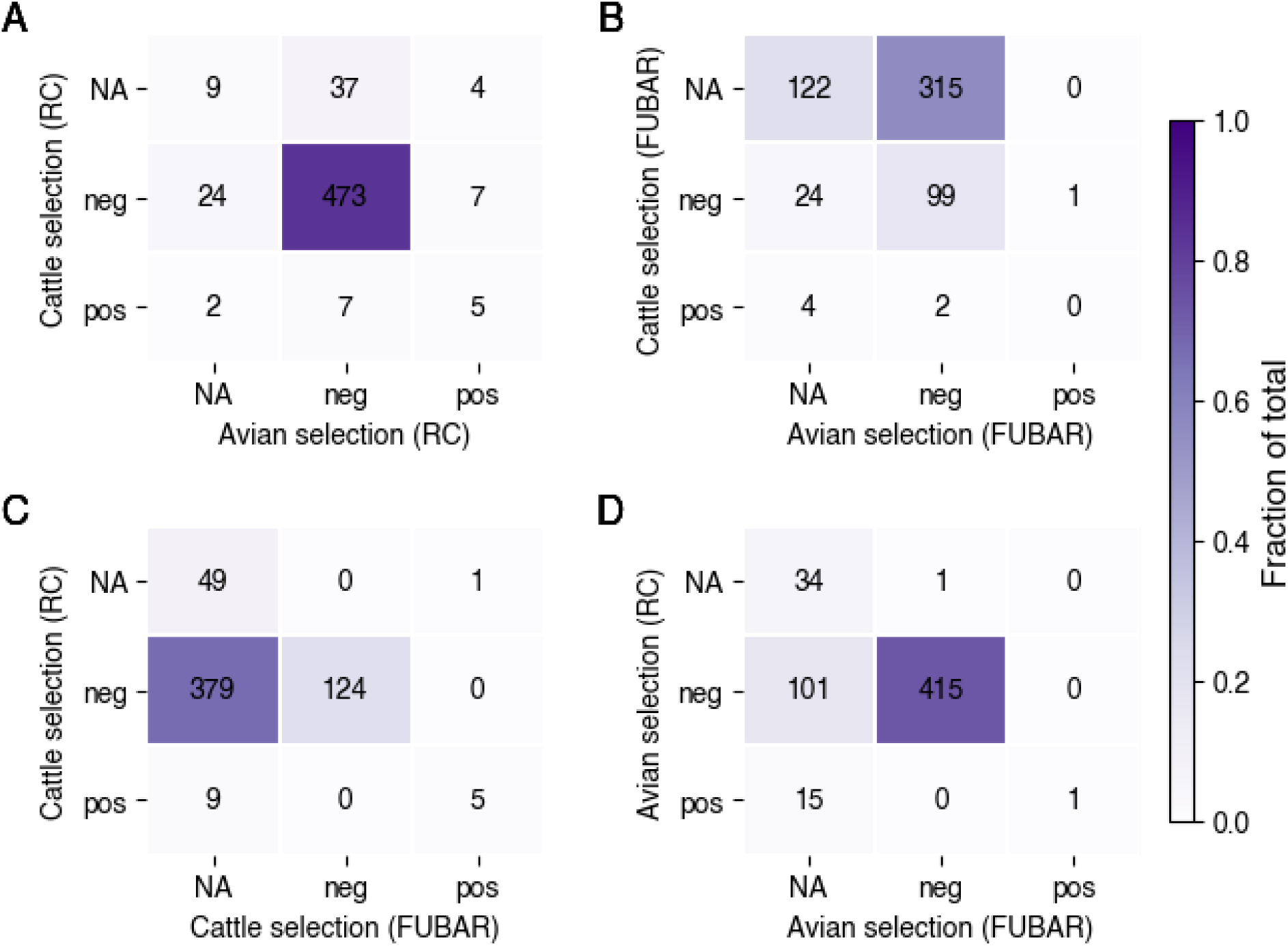
Selection result comparison for the HA gene. **a**–**b**, Comparison of positively, negatively, and neutrally selected for sites in dairy cattle and avian hosts using either robust counting (**a**) or FUBAR (**b**). **c**–**d**, Comparison of robust counting and FUBAR results for sites under selection in (**c**) dairy cattle and (**d**) avian hosts. Stop codons are excluded from FUBAR analyses, resulting in a one-site difference in the total counts between robust counting and FUBAR. RC, robust counting.

**Figure S12.**
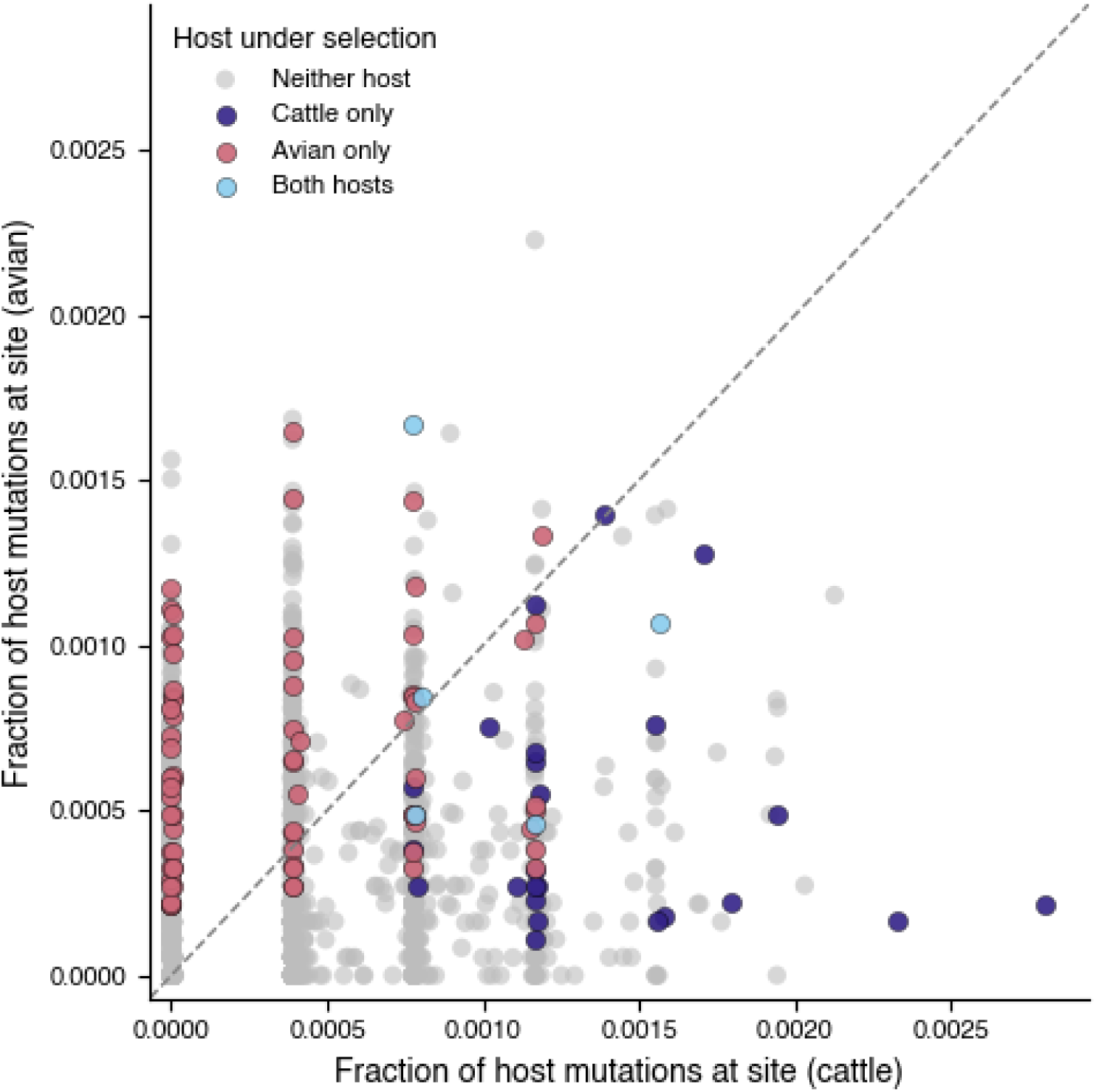
Relationship between substitution frequency and selection among avian and cattle hosts. Points are colored by whether the site is under positive selection and in what host(s), based on robust counting. The fraction of host mutations per site is based on normalization across all segments.

**Figure S13.**
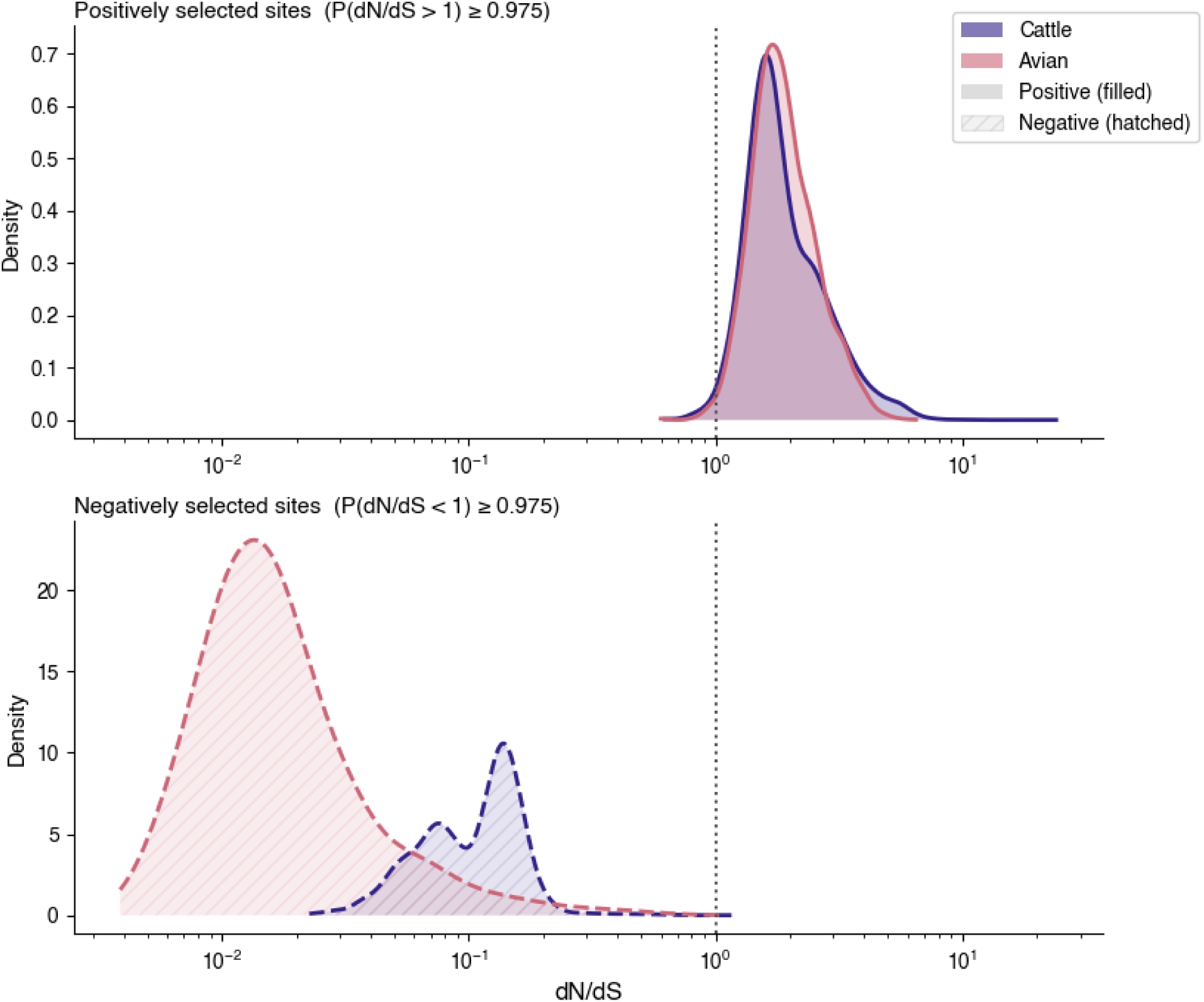
Posterior dN/dS distributions for sites under positive selection (top) and negative selection (bottom) in dairy cattle (purple) and avian hosts (red), estimated using the robust counting framework. Each distribution is a kernel density estimate derived from pooled posterior samples across all classified sites (n = 5,000 total samples per host per category, drawn proportionally across sites). Sites were classified as positively selected if P(dN/dS > 1) ≥ 0.975 and as negatively selected if P(dN/dS < 1) ≥ 0.975 across posterior samples. The dashed vertical line indicates dN/dS = 1. Filled areas correspond to positively selected sites; hatched areas correspond to negatively selected sites.

**Figure S14.**
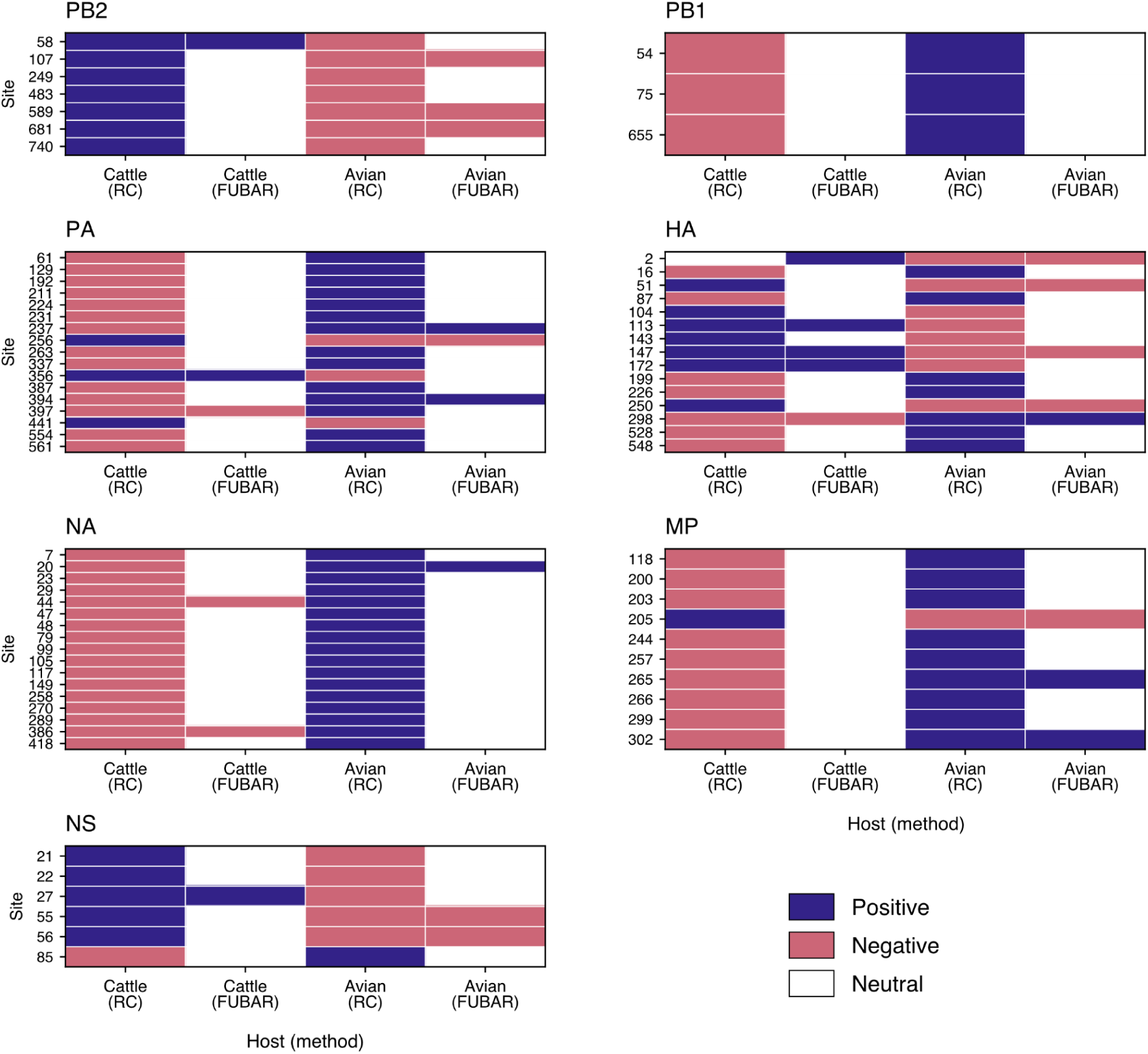
Sites with discordant selection patterns in avian and bovine hosts. Sites that are either positively selected for in dairy cattle and negatively selected for in avian hosts, or vice-versa, when using either robust counting or FUBAR. The NP segment is not shown because it does not have any sites that are either positively selected for in cattle and negatively selected for in birds or negatively selected for in cattle and positively selected for in birds. RC, robust counting.

**Figure S15.**
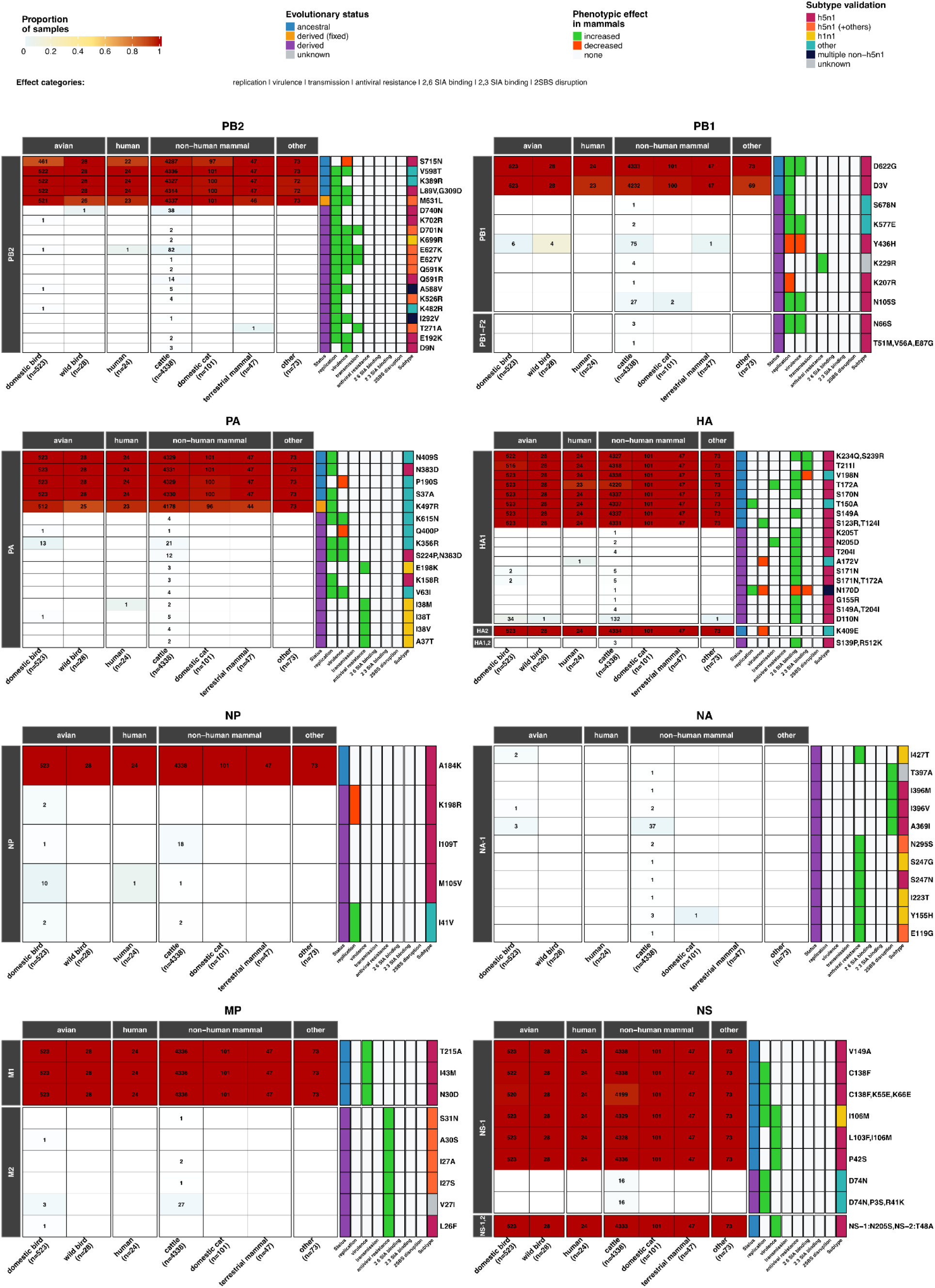
Proportion of characterized phenotypic markers found among B3.13 host groups. Heatmaps for each segment are annotated with: the number of samples, if any, with the respective amino acid substitutions, the phenotypic marker evolutionary status (i.e., whether the marker is ancestral to the emergence of the genotype, derived and subsequently fixed in most of the samples, or derived in a minority of samples), the phenotypic effect of the marker in mammals, and which subtypes the marker has been experimentally validated in. For HA, residues are reported using sequential numbering, including those in the HA2 domain. Figure details and associated citations can be found in Table S6, which uses the sequential HA1-HA2 numbering. Conversions for HA can be found in Table S5.

**Figure S16.**
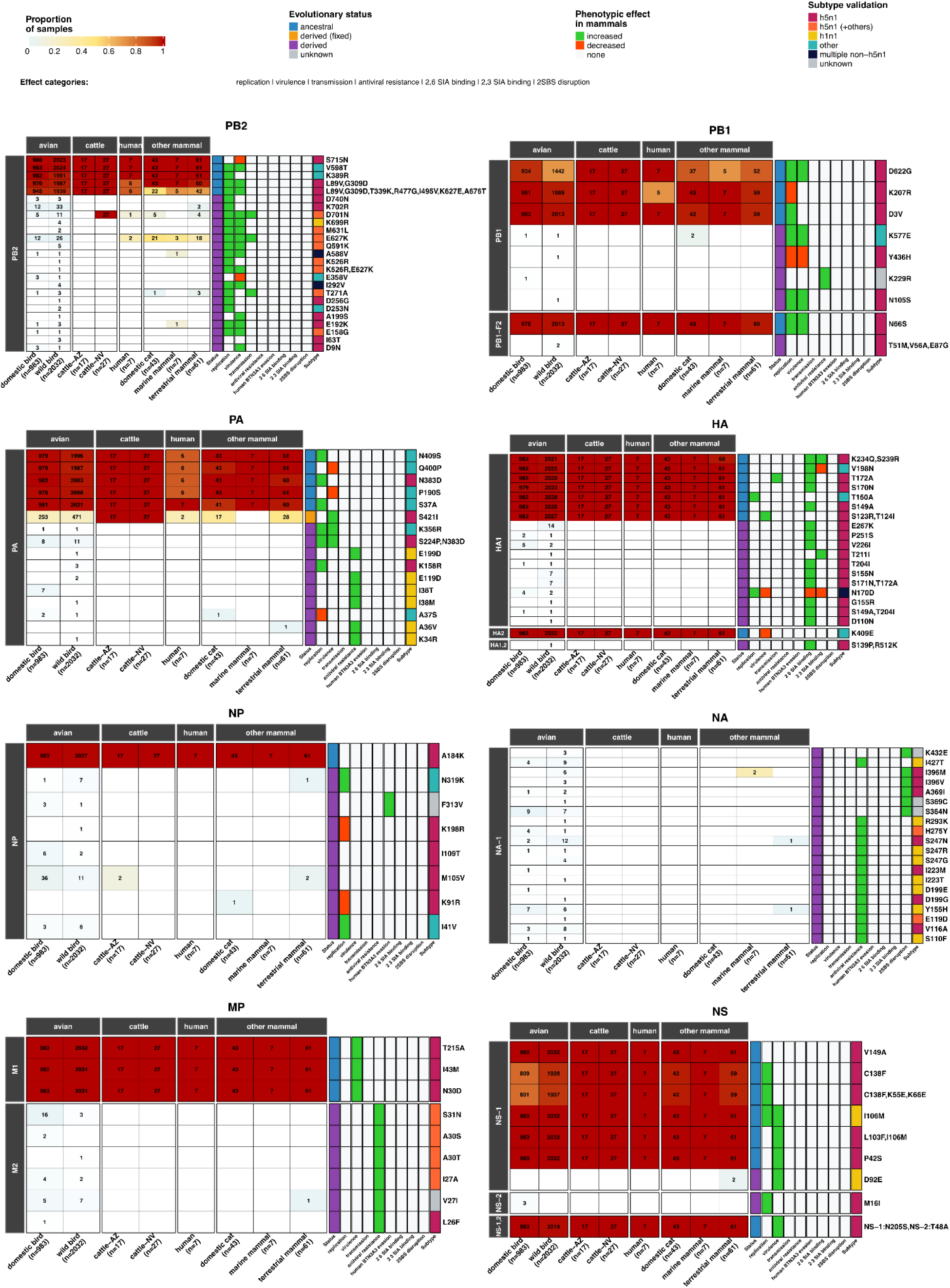
Proportion of characterized phenotypic markers found among D1.1 host groups. Heatmaps for each segment are annotated with: the number of samples, if any, with the respective amino acid substitutions, the phenotypic marker evolutionary status (i.e., whether the marker is ancestral to the emergence of the genotype, derived and subsequently fixed in most of the samples, or derived in a minority of samples), the phenotypic effect of the marker in mammals, and which subtypes the marker has been experimentally validated in. For HA, residues are reported using sequential numbering, including those in the HA2 domain. Figure details and associated citations can be found in Table S6, which uses the sequential HA1-HA2 numbering. Conversions for HA can be found in Table S5.

**Figure S17.**
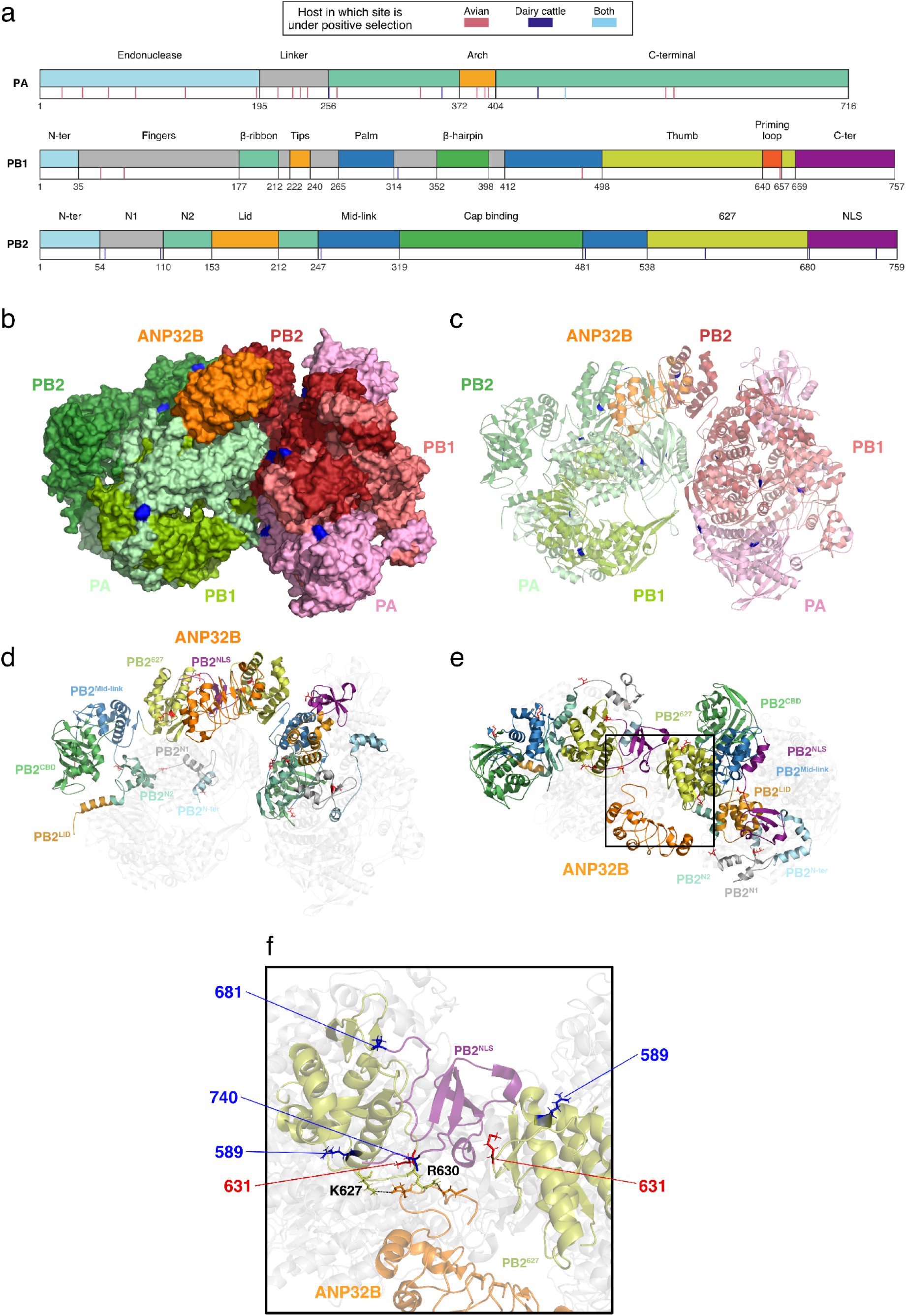
Structural representation of the influenza A virus polymerase complex (FluPolA). **a,** Domain organization of the FluPolA subunits PA, PB1, and PB2, with positively selected sites inferred by robust counting indicated for each subunit. **b,** Surface representation of the FluPolA dimer from the avian H5N1 strain A/turkey/Turkey/1/2005 in complex with human ANP32B (PDB: 8R1J), comprising two polymerase heterotrimers (blue and green) and ANP32B (orange). Sites under positive selection in dairy cattle are shown in red. **c,** Cartoon representation of the FluPolA dimer in complex with human ANP32B. **d,** Cartoon representation of the FluPolA dimer and ANP32B, with only the PB2 subunits of each heterotrimer colored by domain. **e,** Same representation as in (**d**), rotated by 90°. **f,** Close-up view of the PB2 627-domain–ANP32B interface, highlighting PB2 sites under positive selection in dairy cattle (red) and site 631 (blue), as the M631L substitution occurred on the branch leading to the B3.13 dairy cattle outbreak clade. Two residues where PB2 binds to ANP32B are annotated. CBD, cap binding domain. NLS, nuclear localization signal.

**Figure S18.**
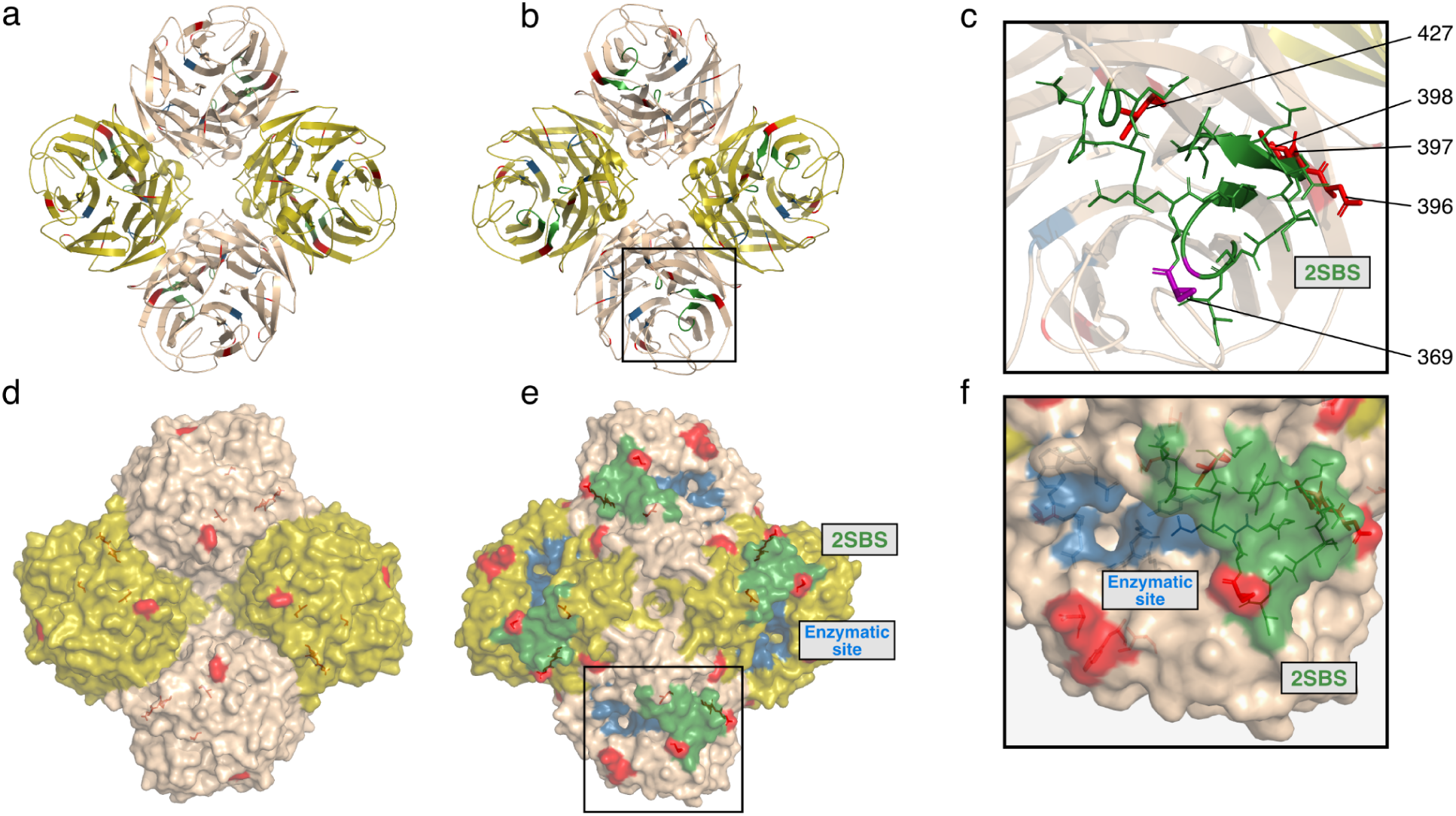
Structural representation of the neuraminidase (NA) head domain. **a–b,** Cartoon representations of the tetrameric NA head domain from the H1N1 strain A/California/04/2009 (PDB: 3NSS), shown in top (**a**) and bottom (**b**) views. **c**, Close-up view of the second sialic acid–binding site (2SBS; green), comprising the 370 loop (residues 366–373), 400 loop (residues 399–404), and 430 loop (residues 430–433), together with nearby key residues (red). A key residue located within the 370 loop is shown in purple. **d–e**, Surface representations of the NA head domain shown from the top (d) and bottom (e) views. **f**, Close-up surface view highlighting the spatial relationship between the second sialic acid–binding site (green) and the enzymatic site (blue). Residues highlighted in red correspond to sites inferred to be neutrally evolving but exhibiting at least three substitutions across the phylogeny, as well as sites classified as virologically important based on phenotypic marker annotations summarized in Fig. S15. Individual protomers are colored in alternating shades to aid visual distinction within the tetramer.

## Supplemental Note 1

### Selection in PB2 and PA

It is thought the influenza virus polymerase is carefully regulated through accumulation of nascent viral proteins and cellular host factors to optimise virus replication, for example by formation of higher order structures (such as ‘dimers of trimers’, two known structures being the ‘symmetrical’ dimer^36^ and the ‘asymmetrical’ dimer^37^), mediated by host ANP32 proteins. However, it is known that mismatches in host factors, such as ANP32, during host switching can lead to host barriers to virus replication that require viral adaptations to overcome^38,39^. We did not infer certain polymerase sites to be under positive selection—likely due to low statistical power concerning non-recurrent but adaptive substitutions—even though particular substitutions at these sites have emerged during circulation in dairy cattle, including PB2 E627K/V, D701N, and Q591R/K, and have been shown to enhance replication in cattle cells through interactions with host ANP32 proteins (Figs. S15, S16, S17; Table S6; refs.^21,28^). These substitutions have been well-described, either individually or together in other mammalian transmission clusters of H5N1^28,40–42^. Several selected for polymerase mutations in cattle lie in potentially functionally relevant interfaces, for example PA 441, PB2 589 and 740 lie proximal to the polymerase-ANP32 binding site (Fig. S17D–F), and could therefore influence interaction with this key host factor^43^. PB2 249 and 589, and PA 441 and 356 all lie on the asymmetric polymerase dimer interface. PA 356 also lies directly on the symmetrical polymerase dimer interface close to other mutations that have been described as allowing influenza virus to overcome host barriers^44^.

### Selection in NA

Influenza NA facilitates virion release through its enzymatic site and, in avian viruses, also contains an adjacent second sialic acid–binding site (2SBS) that has been implicated in mammalian adaptation^45^ and was repeatedly seen in mink farm outbreaks of H5N1^40,46^. In cattle-associated B3.13 viruses, we observed several substitutions in and around the NA 2SBS that show evidence of repeated substitution despite being inferred as neutrally evolving (Figs. S15, S18; Table S3).

### Selection in bovine versus avian hosts

With the exception of PB2, HA, and NS, most gene segments exhibited sites under positive selection predominantly in avian rather than bovine hosts (Tables S3, S4). Positively selected sites in NS were observed in both hosts, whereas sites under positive selection in PB2 were detected exclusively in bovine viruses.

Additionally, a number of sites exhibited opposing selection regimes between hosts, appearing to be under positive selection in one host while being under negative selection in the other host (Fig. S14). In the HA gene, this pattern was observed in both directions, with some sites under positive selection in dairy cattle and negative selection in avian hosts, and other sites under positive selection in avian hosts and negative selection in dairy cattle. In NS and PB2, there were more sites under positive selection in bovine hosts and under negative selection in avian hosts. The remaining gene segments predominantly exhibited sites under positive selection in avian hosts while being under negative selection in bovine hosts.

Although site-specific results inferred using the FUBAR framework were broadly consistent with those obtained from robust counting, FUBAR was generally more conservative in assigning sites to positive or negative selection categories in both hosts. Only two sites in HA—positions 147 and 298—were inferred by FUBAR to be under opposing selective regimes between hosts; at most other sites, FUBAR inferred selection in only one host or neutral evolution in both hosts (Fig. S14).

## Methods

### Dataset compilation

An H5Nx clade 2.3.4.4b dataset for North America, South America and Antarctica was compiled by querying the Global Initiative on Sharing All Influenza Data (GISAID; https://gisaid.org/)^47^ and NCBI Virus^48^ databases for all H5Nx clade 2.3.4.4b samples collected between November 1, 2021 – September 15, 2025 with available genomic sequences for all eight gene segments (accessed on September 15, 2025). Additionally, consensus genomes from raw reads deposited to the NCBI Sequence Read Archive and associated with BioProjects PRJNA1102327, PRJNA1122849, PRJNA1134696, PRJNA1219588, PRJNA1207547, PRJNA980729 were downloaded from the Andersen Lab’s avian-influenza github repository (https://github.com/andersen-lab/avian-influenza). The samples in each dataset were assigned a genotype using the GenoFLU genotyping tool^49^. Sequence datasets for genotypes B3.13 (*n* = 5,134) and D1.1 (*n* = 3,177) were compiled for each gene segment (i.e., PB2, PB1, PA, HA, NP, NA, MP and NS) after removing duplicate sample entries across databases (Table S7).

To assess the molecular evolution of each genome segment before and after the emergence of B3.13 in cattle and place the B3.13 cattle clade in broader phylogenetic context, we compiled background datasets of avian virus sequences for each segment (Table S7). These datasets included sequences for the B3.13 clade 2.3.4.4b Eurasian-derived segments from the ‘ea1’ lineages as defined by GenoFLU (i.e., PA, HA, NA, and MP from genotype A1 and its reassortants) and segments derived from North American low pathogenicity avian influenza viruses or LPAIV (i.e., PB2, PB1, NP, and NS). The sequences for the Eurasian-derived segments were compiled as described for the B3.13 and D1.1 datasets, while only retrieving sequences from clade 2.3.4.4b samples that shared the same GenoFLU lineage for each corresponding segment, were collected from avian hosts, and had complete collection dates (accessed on January 7, 2025). Initial datasets were deduplicated first by accession number, and subsequently, all except one isolate with identical sequences, collection dates, and host metadata were removed using CAMPI (https://github.com/jlcherry/CAMPI). For the LPAIV-derived segments, we queried the GISAID database for all non-H5N1 influenza A samples collected in North America, South America and Antarctica between January 1, 2015 - January 7, 2025 with complete genomes. LPAIV samples were analyzed with GenoFLU and segment sequences that were assigned a shared lineage with a B3.13 segment were retrieved. The resulting background avian dataset for B3.13 included 7709 total avian samples across eight segment datasets: PB2 (*n*= 910), PB1 (*n*= 608), PA (*n*= 5401), HA (*n*= 5555), NP (*n*= 15), NA (*n*= 5363), MP (*n*= 4646) and NS (*n*= 2470).

For each segment, we subsampled the B3.13 background avian virus genome datasets by allowing up to 50 genomes per month while maximizing phylogenetic diversity using Treemmer v0.3 (ref.^50^). For months with fewer than 50 genomes, all available genomes were retained. For October–December 2023, all avian virus genomes were included to maximize diversity, as this period likely corresponds to the emergence of B3.13 in dairy cattle. Phylogenetic outliers were filtered using TreeTime^51^. These datasets excluded all mammalian virus genomes with the exception of a single skunk B3.13 genome, which likely represents a dead-end infection; the branch leading to this virus is inferred to predominantly reflect evolution in the avian reservoir, and it is one of the only B3.13 genomes not associated with the dairy cattle outbreak. For the dairy cattle B3.13 virus genome dataset, we similarly allowed up to 50 genomes per month while maximizing phylogenetic diversity using Treemmer. All human virus genomes nested within the dairy cattle outbreak were included, as there is currently no evidence of human-to-human transmission of B3.13; human infections of B3.13 therefore likely represent sentinel infections rather than sustained human-to-human transmission. All other mammalian virus genomes and all avian virus genomes associated with the dairy cattle outbreak were excluded, as our analyses focus specifically on viral evolution in dairy cattle. All datasets were truncated at May 2025, as we are focused on viral evolution in the first year after emergence.

We constructed A1 alignments to improve phylogenetic resolution, by concatenating the PA, HA, NA, and MP segments of virus genomes present in each of the four individual datasets. This was done for both the avian background and dairy cattle outbreak datasets.

We additionally constructed a B3.13 alignment by concatenating all eight segments of virus genomes present in each of the eight individual datasets.

For the D1.1 avian virus dataset, we first constructed a maximum likelihood phylogeny using all available D1.1 genomes using whole genomes, after which a dataset comprising both closely related sibling lineages and broader phylogenetic diversity was manually selected. All cattle virus genomes were included. As with B3.13, all human virus genomes were retained, and mammalian virus genomes that are immediately sibling to the dairy cattle outbreak clades—and thus could plausibly represent spillovers from cryptic transmission among dairy cattle—were included. All other mammalian virus genomes and all poultry virus genomes associated with the dairy cattle outbreaks were excluded, as our analyses focus on evolution in dairy cattle. No temporal cutoff was applied for D1.1. As there is no documented reassortment within the D1.1 lineage and it was an already established lineage circulating in wild birds, we did not need to construct segment-specific datasets.

At the time of writing, a D1.1 genome from the Wisconsin premise associated with a recent independent spillover (see Figure S7) became available. However, because this sequence was released after the cutoff date for the dairy cattle dataset (15 September 2025) and represents a single genome, it was not included in downstream Bayesian analyses.

### Maximum likelihood phylogenetic analysis of B3.13 and background avian viruses

We used MAPLE v0.7.5 to perform phylogenetic inference on each genome segment, the A1 concatenated alignment composed of the PA, HA, NA, and MP segments, and the whole-genome B3.13 alignment, using a generalized time-reversible (GTR) nucleotide substitution model and the remaining standard settings^52^.

### Bayesian phylogenetic analysis of B3.13 and background avian viruses

We used BEAST X v10.5.0 to perform phylogenetic inference across each segment of H5N1 B3.13, as well as the concatenated alignment of PA-HA-NA-MP^16^. For each of segments and the concatenated alignment, three analyses were performed: an analysis just on the avian virus background dataset (including the single B3.13 skunk virus), an analysis on the dairy cattle outbreak (including the B3.13 human viruses), and an analysis on these two datasets combined.

For each of the analyses of the individual segments for the avian virus background and the dairy cattle outbreak, we used a non-parametric skygrid prior^53^ with 35 gridpoints and a cutoff of 6 years, an uncorrelated relaxed clock^54^, a codon-partitioned model with a GTR substitution model for each codon position, and Hamiltonian Monte Carlo gradient-based sampling for node ages and branch rates^55^. We used robust counting to infer the number of synonymous and nonsynonymous substitutions on branches^15,56^. For each analysis, we first ran one chain of 100 million generations, subsampling every 50 thousand iterations to continuous parameter log and tree files. Convergence and mixing were assessed in Tracer v1.7.2 (ref.^57^), and between 10% and 50% of the chain was discarded as burn-in for each analysis. For analyses where any relevant ESS values were <200, we ran an additional 1–2 chains with the same specifications as the first, discarded the first 10–50% of the chain as burn-in, and combined the chains using LogCombiner. All relevant ESS values for the 1–3 chain analyses were >200.

For the concatenated alignments for the avian virus background and the dairy cattle outbreak, we performed two sets of analyses. We first used the same parameterization as the above analyses for the individual segments, except on the entire A1 concatenated alignment.

We then performed an analysis where the four concatenated segments were treated as separate partitions, with separate clock rates and substitution models, but shared the same tree. The remaining parameterizations were the same as above, except we used standard MCMC operators rather than Hamiltonian Monte Carlo gradient-based sampling.

For each of the analyses of the A1 concatenated alignments, we again ran one chain of 100 million generations, subsampling every 50 thousand iterations to continuous parameter log and tree files. Convergence and mixing were assessed in Tracer, and between 10% and 50% of the chain was discarded as burn-in for each analysis. For analyses where any relevant ESS values were <200, we ran an additional 1–2 chains with the same specifications as the first, discarded the first 10–50% of the chain as burn-in, and combined the chains using LogCombiner. All relevant ESS values for the 1–3 chain analyses were >200.

For the analyses of the combined avian background and dairy cattle outbreak datasets, we used a joint population model, with a constant size population model for the avian background and an exponential population model for the dairy cattle outbreak. We used a codon-partitioned model, except here there were two partitions, with the first and second codon position as one partition and the third codon position as the other partition. We used a GTR+G4 substitution model for each codon partition. For each analysis, we first ran one chain of 100 million generations, subsampling every 50 thousand iterations to continuous parameter log and tree files. Convergence and mixing were assessed in Tracer, and between 10% and 50% of the chain was discarded as burn-in for each analysis. For analyses where any relevant ESS values were <200, we ran an additional 1–2 chains with the same specifications as the first, discarded the first 10–50% of the chain as burn-in, and combined the chains using LogCombiner. All relevant ESS values for the 1–3 chain analyses were >200. The same parameterization was used for the concatenated alignments.

### Maximum likelihood phylogenetic analysis of D1.1

We used MAPLE v0.7.5 to perform phylogenetic inference on a concatenated alignment of all eight segments, using a generalized time-reversible (GTR) nucleotide substitution model and the remaining standard settings.

### Bayesian phylogenetic analysis of D1.1

We used BEAST X to perform phylogenetic inference on the coding regions of each gene segment. Because the dataset included all 8 segments, we performed our analysis in two sequential steps: we first generated an empirical tree distribution and then performed our robust counting inference using the resulting tree distribution.

For the first step of the inference, we treated the eight segments as separate partitions, with a shared tree, clock rates, and codon-partitioned substitution models. We used an uncorrelated relaxed clock and a non-parametric skygrid prior with 25 gridpoints and a cutoff of one year. For the viruses that were sampled in 2024 but do not have complete collection dates, we sampled uniformly from a 3-month window, representing the last 3 months of 2024, as D1.1 was first detected in October 2024. For the viruses sampled in 2025 but do not have complete collection dates, we sampled uniformly from a 2-week window centered on the date 3 weeks prior to the publication date. The viruses sampled in 2025 without complete collection dates were predominantly cattle viruses, but also included several avian and mammalian viruses. We ran 17 chains of 100 million generations, subsampling every 10 thousand iterations to continuous parameter log and tree files. Convergence and mixing were assessed in Tracer, and between 10% of each chain was discarded as burn-in. The 17 chains were then combined and downsampled using LogCombiner, to construct an empirical tree distribution with a total of 1,530 trees. All relevant ESS values were >150.

For the second step of the inference, we used the empirical tree distribution described above, and again treated the eight segments as separate partitions, with a shared tree, the same enforced clock rates, and codon-partitioned substitution models. We used robust counting to infer the number of substitutions on branches and their neutral expectations for each segment. We simulated one MCMC chain of 10 million generations, subsampling every 1,000 iterations to continuous log and tree files. Convergence and mixing were assessed in Tracer, and we discarded the first 10% of the chain as burn-in.

All phylogenetic trees were processed using TreeSwift v1.1.42 and visualized using baltic v0.2.2 (refs.^58,59^).

### Testing for change in intensity of natural selection using RELAX

To explore potential changes in natural selection since the MRCA of H5N1 in cattle, we analyzed maximum likelihood phylogenies from non-overlapping coding regions. For B.3.13, analyzed the concatenated segments PA, HA, NA, and MP in a single analysis due to their phylogenetic congruence; NP, NS, PB1, and PB2 segments were analyzed independently due to their reassortant phylogenies. For D.1.1, all genomic segments were analyzed in a single analysis, due to phylogenetic congruence. We compared the intensity of natural selection between phylogenetic branches representing avian and cattle H5N1 virus using the RELAX^14^ framework in HyPhy v.2.5.1 (ref.^60^). RELAX models heterogeneity in selective pressure across sites and branches by representing the distribution of the nonsynonymous to synonymous substitution rates (ω) as a small number of discrete categories, spanning purifying (ω < 1), neutral (ω ≈ 1), and diversifying (ω > 1) selection. For this hypothesis testing framework, the ω distribution for the background branches (avian H5N1 viruses) and the ω distribution on the test branches (dairy cattle H5N1 outbreak) is modeled as a scaled transformation, parameterized by a single scalar, k. Values of K < 1 indicate a shift of ω values toward neutrality, consistent with relaxation of selection in dairy cattle virus relative to avian virus; values of K > 1 indicate a shift away from neutrality, consistent with intensified selection. Because our aim was to compare the selective environment in dairy cattle with that in wild birds, we excluded the stem branches from the focal comparison. Branches not assigned to either background or test sets, including all branches leading to viruses sampled from mammals other than humans and cattle, were treated as a nuisance class with their own inferred ω distribution.

For each analysis, we modeled site-to-site synonymous rate variation^61^. We also explored the sensitivity of our inference to the number of model ω parameters: 2 versus 3. Model fit was assessed using AIC.

### Identifying sites under diversifying selection using FUBAR

We used FUBAR to identify sites codon that experienced strong diversifying (positive) selection^17^. For each genomic region, we analyzed avian viruses from cattle viruses separately across their section of the maximum likelihood phylogenies. Posterior support of 0.9 (for dN – dS > 0) was used for determining statistical significance.

### Mapping positively selected sites onto protein structures

To visualize the structural context of sites inferred to be under positive selection, we mapped site-specific selection results onto high-resolution protein structures for HA (PDB: 9DWE)^62^, NA (PDB: 3NSS)^63^, and the influenza polymerase complex (PDB: 8R1J; a dimer of heterotrimers bound to human ANP32B)^43^ using PyMOL v3.1.0. Structures were selected based on sequence completeness, resolution, and relevance to circulating H5 viruses.

### Identification of experimentally-validated phenotypic markers

Laboratory-characterized markers related to changes in replication, virulence, transmission, antiviral resistance and other phenotypes during mammalian infections were identified within the complete D1.1 (*n* = 3,177) and B3.13 (*n* = 5,134) nucleotide segment alignments using FluMutGUI v3.2.0 (ref.^64^).

## Supplemental information

**Table S1. Evolutionary rates of B3.13 in bovine and avian hosts.**

**Table S2. RELAX results for B3.13 and D1.1.**

**Table S3. Robust counting results for B3.13.**

**Table S4. FUBAR results for B3.13.**

**Table S5. Influenza HA numbering conversion table.**

**Table S6. Characterized phenotypic markers for substitutions observed in B3.13 and D1.1.**

**Table S7. Influenza genome metadata and accessions.**

## Data and code availability

All genome accessions are available in Table S7. The XML files, code, and Newick trees are available at https://github.com/pekarj/h5n1_dairy_cattle_multiple_emergence.

## Acknowledgements.

P.L. acknowledges support by the Research Foundation - Flanders (‘Fonds voor Wetenschappelijk Onderzoek - Vlaanderen’ G051322N and G051323N). T.P. is funded by the UK Medical Research Council/Department for Environment, Food and Rural Affairs (DEFRA) FluTrailMap-One Health consortium (MR/Y03368X/1), the Biotechnology and Biological Sciences Research Council (BBSRC)/DEFRA ‘FluTrailMap’ consortium (BB/Y007298/1), and the BBSRC via the Pirbright Institute’s Strategic Program Grants (ISPGs) (BBS/E/PI/230002A and BBS/E/PI/230002B). This project has received funding from the Research Council of Lithuania, Lithuania under the EMBO Installation Grant programme under grant No 5305 (G.D.). D.H.G. was supported by Academy of Medical Sciences Springboard Grant 1049. This work is supported by the Centers of Excellence for Influenza Research and Response, National Institute of Allergy and Infectious Diseases, National Institutes of Health (NIH), Department of Health and Human Services, under contracts 75N93021C00015 (M.W.) and 75N93021C00014 (M.I.N.), and by the Division of Intramural Research of the US National Library of Medicine at the NIH (M.I.N.). This work is also partially supported through NIH grants U19 AI135995 (K.G.A.), R01 AI153044 (M.A.S) and R01 AI192139 (J.O.W.). M.U.G.K. acknowledges funding from The Rockefeller Foundation (PC-2022-POP-005, also J.E.P. & A.R.), Health AI Programme from Google.org, the Oxford Martin School Programmes in Pandemic Genomics & Digital Pandemic Preparedness, European Union’s Horizon Europe programme projects MOOD (#874850) and E4Warning (#101086640), Wellcome Trust grants 303666/Z/23/Z, 226052/Z/22/Z & 228186/Z/23/Z, the United Kingdom Research and Innovation (#APP8583), the Medical Research Foundation (MRF-RG-ICCH-2022-100069), UK International Development (301542-403), the Bill & Melinda Gates Foundation grants (INV-063472, INV-090281) and Novo Nordisk Foundation (NNF24OC0094346). The contents of this publication are the sole responsibility of the authors and do not necessarily reflect the views of the European Commission or the other funders. The content does not necessarily reflect the views or policies of the Department of Health and Human Services, nor imply endorsement by the U.S. Government. We gratefully acknowledge the authors from the originating laboratories and the submitting laboratories, who generated and shared through GISAID the viral genomic sequences and metadata on which this research is based (Table S7).

## References

1. Hu, X., Saxena, A., Magstadt, D.R., Gauger, P.C., Burrough, E., Zhang, J., Siepker, C., Mainenti, M., Gorden, P.J., Plummer, P., et al. (2024). Highly Pathogenic Avian Influenza A (H5N1) clade 2.3.4.4b Virus detected in dairy cattle. bioRxiv, 2024.04.16.588916. 10.1101/2024.04.16.588916.

2. Burrough, E.R., Magstadt, D.R., Petersen, B., Timmermans, S.J., Gauger, P.C., Zhang, J., Siepker, C., Mainenti, M., Li, G., Thompson, A.C., et al. Early Release - Highly Pathogenic Avian Influenza A(H5N1) Clade 2.3.4.4b Virus Infection in Domestic Dairy Cattle and Cats, United States, 2024 - Volume 30, Number 7—July 2024 - Emerging Infectious Diseases journal - CDC. 10.3201/eid3007.240508.

3. Nguyen, T.-Q., Hutter, C., Markin, A., Thomas, M.N., Lantz, K., Killian, M.L., Janzen, G.M., Vijendran, S., Wagle, S., Inderski, B., et al. (2024). Emergence and interstate spread of highly pathogenic avian influenza A(H5N1) in dairy cattle. bioRxiv, 2024.05.01.591751. 10.1101/2024.05.01.591751.

4. Uyeki, T.M., Milton, S., Abdul Hamid, C., Reinoso Webb, C., Presley, S.M., Shetty, V., Rollo, S.N., Martinez, D.L., Rai, S., Gonzales, E.R., et al. (2024). Highly pathogenic avian influenza A(H5N1) virus infection in a dairy farm worker. N. Engl. J. Med. 390, 2028–2029.

5. Pekar, J.E., Crespo-Bellido, A., Lemey, P., Bowman, A.S., Peacock, T.P., Ochoa, J.N., Rambaut, A., Pybus, O.G., Worobey, M., and Nelson, M.I. (2026). Can H5N1 avian influenza in dairy cattle be contained in the US? Cell. 10.1016/j.cell.2025.12.033.

6. National Milk Testing Strategy (2024). Animal and Plant Health Inspection Service. https://www.aphis.usda.gov/livestock-poultry-disease/avian/avian-influenza/hpai-detections/livestock/nmts.

7. CDC (2025). Genetic Sequences of Highly Pathogenic Avian Influenza A(H5N1) Viruses Identified in a Person in Louisiana. Avian Influenza (Bird Flu). https://www.cdc.gov/bird-flu/spotlights/h5n1-response-12232024.html.

8. Jassem, A.N., Roberts, A., Tyson, J., Zlosnik, J.E.A., Russell, S.L., Caleta, J.M., Eckbo, E.J., Gao, R., Chestley, T., Grant, J., et al. (2025). Critical illness in an adolescent with influenza A(H5N1) virus infection. N. Engl. J. Med. 392, 927–929.

9. Environmental Health Services Division Newsletter - Week of January 5, 2026 (2024). Central Nevada Health District. https://www.centralnevadahd.org/community-alerts-2/.

10. Campbell, A.J., Brizuela, K., and Lakdawala, S.S. (2025). mGem: Transmission and exposure risks of dairy cow H5N1 influenza virus. MBio 16, e0294424.

11. Nguyen, T.-Q., Hutter, C.R., Markin, A., Thomas, M., Lantz, K., Killian, M.L., Janzen, G.M., Vijendran, S., Wagle, S., Inderski, B., et al. (2025). Emergence and interstate spread of highly pathogenic avian influenza A(H5N1) in dairy cattle in the United States. Science 388, eadq0900.

12. Worobey, M., Han, G.-Z., and Rambaut, A. (2014). A synchronized global sweep of the internal genes of modern avian influenza virus. Nature 508, 254–257.

13. Crespo-Bellido, A., Trovão, N.S., Maksiaev, A., Baele, G., Dellicour, S., and Nelson, M.I. (2025). Emergence of D1.1 reassortant H5N1 avian influenza viruses in North America. bioRxiv, 2025.12.19.695329. 10.64898/2025.12.19.695329.

14. Wertheim, J.O., Murrell, B., Smith, M.D., Kosakovsky Pond, S.L., and Scheffler, K. (2015). RELAX: detecting relaxed selection in a phylogenetic framework. Mol. Biol. Evol. 32, 820–832.

15. Lemey, P., Minin, V.N., Bielejec, F., Kosakovsky Pond, S.L., and Suchard, M.A. (2012). A counting renaissance: combining stochastic mapping and empirical Bayes to quickly detect amino acid sites under positive selection. Bioinformatics 28, 3248–3256.

16. Baele, G., Ji, X., Hassler, G.W., McCrone, J.T., Shao, Y., Zhang, Z., Holbrook, A.J., Lemey, P., Drummond, A.J., Rambaut, A., et al. (2025). BEAST X for Bayesian phylogenetic, phylogeographic and phylodynamic inference. Nat. Methods 22, 1653–1656.

17. Murrell, B., Moola, S., Mabona, A., Weighill, T., Sheward, D., Kosakovsky Pond, S.L., and Scheffler, K. (2013). FUBAR: a fast, unconstrained bayesian approximation for inferring selection. Mol. Biol. Evol. 30, 1196–1205.

18. Miyakawa, K., Ota, M., Sano, K., Momose, F., Kishida, N., Arita, T., Suzuki, Y., Shirakura, M., Asanuma, H., Watanabe, S., et al. (2025). Emergence of antigenic variants in bovine H5N1 influenza viruses. J. Med. Virol. 97, e70394.

19. Dadonaite, B., Ahn, J.J., Ort, J.T., Yu, J., Furey, C., Dosey, A., Hannon, W.W., Vincent Baker, A.L., Webby, R.J., King, N.P., et al. (2024). Deep mutational scanning of H5 hemagglutinin to inform influenza virus surveillance. PLoS Biol. 22, e3002916.

20. Peacock, T.P., Sheppard, C.M., Lister, M.G., Staller, E., Frise, R., Swann, O.C., Goldhill, D.H., Long, J.S., and Barclay, W.S. (2023). Mammalian ANP32A and ANP32B proteins drive differential polymerase adaptations in avian influenza virus. J. Virol. 97, e0021323.

21. Dholakia, V., Quantrill, J.L., Richardson, S., Pankaew, N., Brown, M.D., Yang, J., Capelastegui, F., Masonou, T., Case, K.-M., Ajeian, J., et al. (2025). Polymerase mutations underlie early adaptation of H5N1 influenza virus to dairy cattle and other mammals. Microbiology.

22. Preliminary report on genomic epidemiology of the 2024 H5N1 influenza A virus outbreak in U.S. cattle (Part 1 of 2) (2024). Virological. https://virological.org/t/preliminary-report-on-genomic-epidemiology-of-the-2024-h5n1-influenza-a-virus-outbreak-in-u-s-cattle-part-1-of-2/970.

23. Update: Genetic Sequencing Results for Wisconsin Dairy Herd Detection of Highly Pathogenic Avian Influenza (2025). Animal and Plant Health Inspection Service. https://www.aphis.usda.gov/news/agency-announcements/update-genetic-sequencing-results-wisconsin-dairy-herd-detection-highly.

24. Buhnerkempe, M.G., Tildesley, M.J., Lindström, T., Grear, D.A., Portacci, K., Miller, R.S., Lombard, J.E., Werkman, M., Keeling, M.J., Wennergren, U., et al. (2014). The impact of movements and animal density on continental scale cattle disease outbreaks in the United States. PLoS One 9, e91724.

25. Sellman, S., Beck-Johnson, L.M., Hallman, C., Miller, R.S., Bonner, K.A.O., Portacci, K., Webb, C.T., and Lindström, T. (2022). Modeling U.S. cattle movements until the cows come home: Who ships to whom and how many? Comput. Electron. Agric. 203, 107483.

26. Detection of avian flu antibodies in Dutch dairy cow: ECDC risk assessment remains unchanged (2026). European Centre for Disease Prevention and Control. https://www.ecdc.europa.eu/en/publications-data/detection-avian-flu-antibodies-dutch-dairy-cow-ecdc-risk-assessment-remains.

27. Gu, C., Maemura, T., Guan, L., Eisfeld, A.J., Biswas, A., Kiso, M., Uraki, R., Ito, M., Trifkovic, S., Wang, T., et al. (2024). A human isolate of bovine H5N1 is transmissible and lethal in animal models. Nature 636, 711–718.

28. Zhang, L., Lai, Y., Cui, Y., Yang, Q., Shao, Y., Ding, S., Wang, H., Wang, L., Gao, G.F., and Deng, T. (2025). Emergence of mammalian-adaptive PB2 mutations enhances polymerase activity and pathogenicity of cattle-derived H5N1 influenza A virus. Nat. Commun., 1–15.

29. Uhart, M.M., Vanstreels, R.E.T., Nelson, M.I., Olivera, V., Campagna, J., Zavattieri, V., Lemey, P., Campagna, C., Falabella, V., and Rimondi, A. (2024). Epidemiological data of an influenza A/H5N1 outbreak in elephant seals in Argentina indicates mammal-to-mammal transmission. Nat. Commun. 15, 9516.

30. Dunham, E.J., Dugan, V.G., Kaser, E.K., Perkins, S.E., Brown, I.H., Holmes, E.C., and Taubenberger, J.K. (2009). Different evolutionary trajectories of European avian-like and classical swine H1N1 influenza A viruses. J. Virol. 83, 5485–5494.

31. Havens, J.L., Kosakovsky Pond, S.L., Zehr, J.D., Pekar, J.E., Parker, E., Worobey, M., Andersen, K.G., and Wertheim, J.O. (2026). Dynamics of natural selection preceding human viral epidemics and pandemics. Cell 0. 10.1016/j.cell.2026.02.006.

32. Bhatt, S., Lam, T.T., Lycett, S.J., Leigh Brown, A.J., Bowden, T.A., Holmes, E.C., Guan, Y., Wood, J.L.N., Brown, I.H., Kellam, P., et al. (2013). The evolutionary dynamics of influenza A virus adaptation to mammalian hosts. Philos. Trans. R. Soc. Lond. B Biol. Sci. 368, 20120382.

33. Li, B., Raghwani, J., Hill, S.C., François, S., Lefrancq, N., Liang, Y., Wang, Z., Dong, L., Lemey, P., Pybus, O.G., et al. (2025). Association of poultry vaccination with interspecies transmission and molecular evolution of H5 subtype avian influenza virus. Sci. Adv. 11, eado9140.

34. https://www.aphis.usda.gov/sites/default/files/dairy-cattle-hpai-tech-brief.pdf.

35. https://www.nmpf.org/wp-content/uploads/2025/01/NMPF_BTM-Sample-Logistics-for-H5N1-2.pdf.

36. Fan, H., Walker, A.P., Carrique, L., Keown, J.R., Serna Martin, I., Karia, D., Sharps, J., Hengrung, N., Pardon, E., Steyaert, J., et al. (2019). Structures of influenza A virus RNA polymerase offer insight into viral genome replication. Nature 573, 287–290.

37. Carrique, L., Fan, H., Walker, A.P., Keown, J.R., Sharps, J., Staller, E., Barclay, W.S., Fodor, E., and Grimes, J.M. (2020). Host ANP32A mediates the assembly of the influenza virus replicase. Nature 587, 638–643.

38. Long, J.S., Giotis, E.S., Moncorgé, O., Frise, R., Mistry, B., James, J., Morisson, M., Iqbal, M., Vignal, A., Skinner, M.A., et al. (2016). Species difference in ANP32A underlies influenza A virus polymerase host restriction. Nature 529, 101–104.

39. Sheppard, C.M., Goldhill, D.H., Swann, O.C., Staller, E., Penn, R., Platt, O.K., Sukhova, K., Baillon, L., Frise, R., Peacock, T.P., et al. (2023). An Influenza A virus can evolve to use human ANP32E through altering polymerase dimerization. Nat. Commun. 14, 6135.

40. Kareinen, L., Tammiranta, N., Kauppinen, A., Zecchin, B., Pastori, A., Monne, I., Terregino, C., Giussani, E., Kaarto, R., Karkamo, V., et al. (2024). Highly pathogenic avian influenza A(H5N1) virus infections on fur farms connected to mass mortalities of black-headed gulls, Finland, July to October 2023. Euro Surveill. 29. 10.2807/1560-7917.ES.2024.29.25.2400063.

41. Tomás, G., Marandino, A., Panzera, Y., Rodríguez, S., Wallau, G.L., Dezordi, F.Z., Pérez, R., Bassetti, L., Negro, R., Williman, J., et al. (2024). Highly pathogenic avian influenza H5N1 virus infections in pinnipeds and seabirds in Uruguay: Implications for bird-mammal transmission in South America. Virus Evol. 10, veae031.

42. Mollett, B.C., Lynton-Jenkins, J.G., Richardson, S., Quantrill, J.L., Byrne, A.M.P., Harvey, R., Adams, L., Proust, A., Brown, M.D., Yang, J., et al. (2025). Disease ecology and zoonotic risk of clade 2.3.4.4b H5N1 high pathogenicity avian influenza in the sub-Antarctic region. Research Square. 10.21203/rs.3.rs-8029950/v1.

43. Staller, E., Carrique, L., Swann, O.C., Fan, H., Keown, J.R., Sheppard, C.M., Barclay, W.S., Grimes, J.M., and Fodor, E. (2024). Structures of H5N1 influenza polymerase with ANP32B reveal mechanisms of genome replication and host adaptation. Nat. Commun. 15, 4123.

44. Idoko-Akoh, A., Goldhill, D.H., Sheppard, C.M., Bialy, D., Quantrill, J.L., Sukhova, K., Brown, J.C., Richardson, S., Campbell, C., Taylor, L., et al. (2023). Creating resistance to avian influenza infection through genome editing of the ANP32 gene family. Nat. Commun. 14, 6136.

45. Du, W., de Vries, E., van Kuppeveld, F.J.M., Matrosovich, M., and de Haan, C.A.M. (2021). Second sialic acid-binding site of influenza A virus neuraminidase: binding receptors for efficient release. FEBS J. 288, 5598–5612.

46. de Vries, E., and de Haan, C.A. (2023). Letter to the editor: Highly pathogenic influenza A(H5N1) viruses in farmed mink outbreak contain a disrupted second sialic acid binding site in neuraminidase, similar to human influenza A viruses. Euro Surveill. 28. 10.2807/1560-7917.ES.2023.28.7.2300085.

47. Shu, Y., and McCauley, J. (2017). GISAID: Global initiative on sharing all influenza data - from vision to reality. Euro Surveill. 22. 10.2807/1560-7917.ES.2017.22.13.30494.

48. Hatcher, E.L., Zhdanov, S.A., Bao, Y., Blinkova, O., Nawrocki, E.P., Ostapchuck, Y., Schäffer, A.A., and Brister, J.R. (2017). Virus Variation Resource - improved response to emergent viral outbreaks. Nucleic Acids Res. 45, D482–D490.

49. Youk, S., Torchetti, M.K., Lantz, K., Lenoch, J.B., Killian, M.L., Leyson, C., Bevins, S.N., Dilione, K., Ip, H.S., Stallknecht, D.E., et al. (2023). H5N1 highly pathogenic avian influenza clade 2.3.4.4b in wild and domestic birds: Introductions into the United States and reassortments, December 2021-April 2022. Virology 587, 109860.

50. Menardo, F., Loiseau, C., Brites, D., Coscolla, M., Gygli, S.M., Rutaihwa, L.K., Trauner, A., Beisel, C., Borrell, S., and Gagneux, S. (2018). Treemmer: a tool to reduce large phylogenetic datasets with minimal loss of diversity. BMC Bioinformatics 19, 164.

51. Sagulenko, P., Puller, V., and Neher, R.A. (2018). TreeTime: Maximum-likelihood phylodynamic analysis. Virus Evol. 4, vex042.

52. De Maio, N., Kalaghatgi, P., Turakhia, Y., Corbett-Detig, R., Minh, B.Q., and Goldman, N. (2023). Maximum likelihood pandemic-scale phylogenetics. Nat. Genet. 55, 746–752.

53. Gill, M.S., Lemey, P., Faria, N.R., Rambaut, A., Shapiro, B., and Suchard, M.A. (2013). Improving Bayesian population dynamics inference: a coalescent-based model for multiple loci. Mol. Biol. Evol. 30, 713–724.

54. Drummond, A.J., Ho, S.Y.W., Phillips, M.J., and Rambaut, A. (2006). Relaxed phylogenetics and dating with confidence. PLoS Biol. 4, e88.

55. Baele, G., Gill, M.S., Lemey, P., and Suchard, M.A. (2020). Hamiltonian Monte Carlo sampling to estimate past population dynamics using the skygrid coalescent model in a Bayesian phylogenetics framework. Wellcome Open Res. 5, 53.

56. O’Brien, J.D., Minin, V.N., and Suchard, M.A. (2009). Learning to count: robust estimates for labeled distances between molecular sequences. Mol. Biol. Evol. 26, 801–814.

57. Rambaut, A., Drummond, A.J., Xie, D., Baele, G., and Suchard, M.A. (2018). Posterior summarization in Bayesian phylogenetics using tracer 1.7. Syst. Biol. 67, 901–904.

58. Moshiri, N. (2020). TreeSwift: A massively scalable Python tree package. SoftwareX 11, 100436.

59. Dudas, G. baltic: baltic - backronymed adaptable lightweight tree import code for molecular phylogeny manipulation, analysis and visualisation. Development is back on the evogytis/baltic branch (i.e. here) (Github).

60. Kosakovsky Pond, S.L., Poon, A.F.Y., Velazquez, R., Weaver, S., Hepler, N.L., Murrell, B., Shank, S.D., Magalis, B.R., Bouvier, D., Nekrutenko, A., et al. (2020). HyPhy 2.5-A customizable platform for evolutionary hypothesis testing using PHYlogenies. Mol. Biol. Evol. 37, 295–299.

61. Wisotsky, S.R., Kosakovsky Pond, S.L., Shank, S.D., and Muse, S.V. (2020). Synonymous site-to-site substitution rate variation dramatically inflates false positive rates of selection analyses: Ignore at your own peril. Mol. Biol. Evol. 37, 2430–2439.

62. Good, M.R., Fernández-Quintero, M.L., Ji, W., Rodriguez, A.J., Han, J., Ward, A.B., and Guthmiller, J.J. (2024). A single mutation in dairy cow-associated H5N1 viruses increases receptor binding breadth. Nat. Commun. 15, 10768.

63. Li, Q., Qi, J., Zhang, W., Vavricka, C.J., Shi, Y., Wei, J., Feng, E., Shen, J., Chen, J., Liu, D., et al. (2010). The 2009 pandemic H1N1 neuraminidase N1 lacks the 150-cavity in its active site. Nat. Struct. Mol. Biol. 17, 1266–1268.

64. Giussani, E., Sartori, A., Salomoni, A., Cavicchio, L., de Battisti, C., Pastori, A., Varotto, M., Zecchin, B., Hughes, J., Monne, I., et al. (2025). FluMut: a tool for mutation surveillance in highly pathogenic H5N1 genomes. Virus Evolution 11. 10.1093/ve/veaf011.

